# Anatomically-based skeleton kinetics and pose estimation in freely-moving rodents

**DOI:** 10.1101/2021.11.03.466906

**Authors:** Arne Monsees, Kay-Michael Voit, Damian J. Wallace, Juergen Sawinski, Edyta Leks, Klaus Scheffler, Jakob H. Macke, Jason N. D. Kerr

## Abstract

Forming a complete picture of the relationship between neural activity and body kinetics requires quantification of skeletal joint biomechanics during behavior. However, without detailed knowledge of the underlying skeletal motion, inferring joint kinetics from surface tracking approaches is difficult, especially for animals where the relationship between surface anatomy and skeleton changes during motion. Here we developed a videography-based method enabling detailed three-dimensional kinetic quantification of an anatomically defined skeleton in untethered freely-behaving animals. This skeleton-based model has been constrained by anatomical principles and joint motion limits and provided skeletal pose estimates for a range of rodent sizes, even when limbs were occluded. Model-inferred joint kinetics for both gait and gap-crossing behaviors were verified by direct measurement of limb placement, showing that complex decision-making behaviors can be accurately reconstructed at the level of skeletal kinetics using our anatomically constrained model.

## Introduction

The relationship between neural activity patterns and body motion is complex as neuronal activity patterns are dependent on factors such as the intended behavioral outcome^1^, task familiarity^2^, changes in the environment but also exact limb trajectories^3–5^ and motion kinetics^6^. Much of the motion kinematic data forming our view of the sensorimotor control of movement was collected during short behavioral epochs where the animal was in various forms of restraint^3–8^, but how these findings relate to kinematics during free behavior, where the relationship between the environment and body motion is continuously changing, is largely unknown^9–11^. While there have been methodological advances recently enabling detailed neural population activity recordings^12–15^ and surface tracking of an animal’s body^16–24^, a major challenge still remains for generating detailed kinetics of individual body parts, such as limbs, and how they interact with the environment during free behavior^18,25^. This poses an especially difficult problem as limb motions involving muscles, bones and joints are biomechanically complex given their three-dimensional (3D) translational and rotational co-dependencies^26,27^.

More recently, advances in the development of machine learning approaches have enabled limb tracking in both freely-moving^23^ and head-restrained insects^28^ as the limb exoskeleton not only provides joint angle limits and hard limits of limb position, but can be tracked as a surface feature during behavior. When studying vertebrates, like rats, the entire skeleton is occluded by the animal’s fur and inferring bone positions and calculating joint kinetics becomes more complicated since the spatial relationship between skeleton and overlying soft tissues are less apparent^29,30^. Despite this limitation, recent approaches have extended two-dimensional surface tracking methods^21,23,24^ to include 3D pose reconstructions^31^ using a multi-camera cross-validation approach and hand-marked ground-truth data sets^19^ allowing general kinematic representation of animal behaviors and poses for multiple species^32^. Extending these approaches to obtain the skeleton kinetics relies on knowledge of the skeleton anatomy and biomechanics as well as motion restrictions of joints^27^ because animal poses are limited by both bone lengths and joint angle limits. Here, we developed an anatomically constrained skeleton model incorporating mechanistic knowledge of bone locations, anatomical limits of bone rotations, and temporal constraints to track 3D joint positions and their kinetics in freely moving rats. Together the fully constrained skeleton enabled the reconstruction of skeleton poses and kinetic quantification during gap-crossing tasks and throughout spontaneous behavioral sequences.

## Results

### Constraining the model

Here we tracked 3D joint positions and their kinetics in freely moving rats (Fig. 1a) over a large size range (N = 6 animals, average weight: 321 g, range: 71-735 g) using videography and a anatomically constrained skeleton model (ACM) incorporating mechanistic knowledge of bone locations, anatomical limits of bone rotations, and temporal constraints. Together, both the temporal and anatomical constraints, i.e. the fully constrained ACM, enabled the reconstruction of skeleton poses of behaving animals with single joint precision (Fig. 1b) as well as smooth limb and joint transitions during gait cycles (Fig. 1c) allowing the quantification joint kinetics and spatial positions of the limbs throughout behavioral sequences. At the core of this approach was a generalized rat skeleton based on rat bone anatomy^33^ (Fig. 1d) modeled as a mathematical graph with vertices representing individual joints and edges representing bones (Fig. 1c, see methods for details and Supplementary Fig. 1). For example, a single edge was used to represent the animal’s head, the spinal column was approximated using four edges based on cervical, thoracic and lumbar sections of the column as well as the sacrum^33^ and the tail by five edges (Fig. 1d, Supplementary Fig. 1). To constrain the model we applied angle limits for each joint based on measured rotations^34^ (Fig. 1e, see Methods “Constraining poses based on joint angle limits”) as well as anatomical constraints based on measured relationships between bone lengths and animal weight and size^35^ (see Methods “Constraining bone lengths based on allometry”). Finally, as vertebrates are symmetrical around the mid-sagittal plane we applied a further anatomical constraint to ensure symmetry for bone lengths and surface marker locations (Supplementary Fig. 3). Together, this approach established a unique skeleton model for each animal. To generate probabilistic estimates of 3D joint positions and provide temporal constraints, we implemented a temporal unscented Rauch-Tung-Striebel (RTS) smoother^36,37^, an extension of a Kalman filter^38^, which is suitable for nonlinear dynamics models and also incorporates information from future marker locations (see Supplementary Methods “Probabilistic pose estimation” for details). Parameters of the smoother were learned via the expectation-maximization (EM) algorithm^39^, by iteratively fitting poses of the entire behavioral sequence (see Methods “Performing probabilistic pose reconstruction”).

**Fig. 1.**
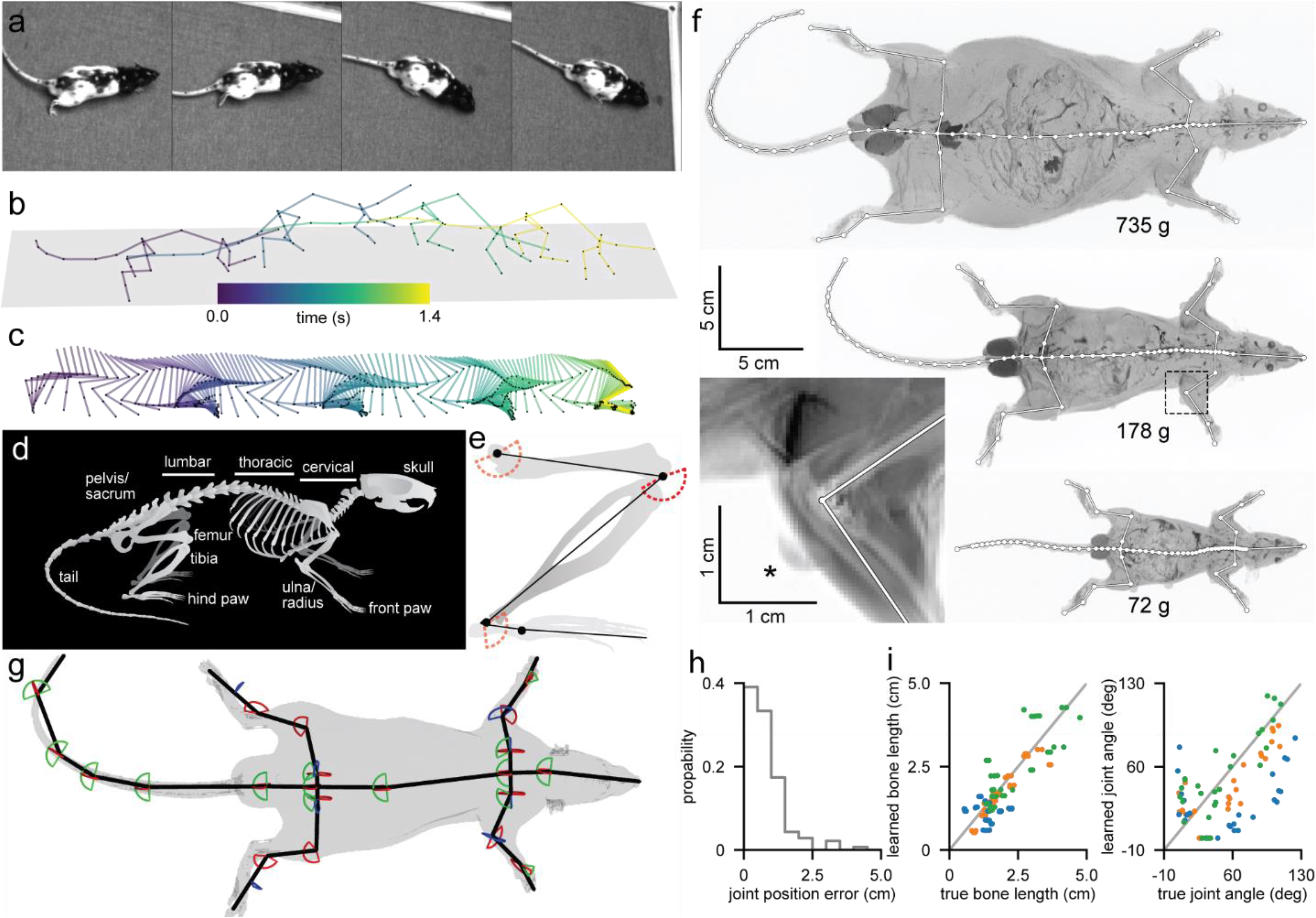
Leaning anatomically informed skeleton models allows for accurate 3D pose reconstruction during free behavior. **a**, Four recorded frames from an overhead camera showing a freely moving rat with painted surface labels. **b**, Reconstructed animal poses of the entire skeleton during gait. Poses correspond to the images shown in a. **c**, Enlargement of the reconstructed right hind limb during the sequence shown in b. **d**, Schematic image of a rat skeleton showing anatomical landmarks. **e**, Schematic image of a hind limb with modeled bones (black lines) and joints (black dots) as well as enforced joint angle limits for flexion/extension (red dashed lines). **f**, MRI scans of three differently sized animals (maximum projection) and an enlargement of a right elbow joint (lower left, mean projection, same area as in left dashed box) with manually labeled bone (white lines) and joint (white dots) positions. Note visible MRI surface marker (asterisk). **g**, 3D representation of a rat’s MRI scan showing the animal’s surface (gray) and the aligned skeleton model (black lines) and joint angle limits for flexion/extension (red lines), abduction/adduction (green lines) and internal/external rotation (blue lines). **h**, Probability histogram of the joint position error. **i**, Learned bone lengths (left) and joint angles (right) compared to MRI bone lengths and joint angles (N = 6 animals). Colors represent small (blue), medium (orange) and large (green) animal sizes (blue: 71 g & 72 g, orange: 174 g & 178 g, green: 699 g & 735 g).

### Learning the skeleton

To relate the animal’s surface to the underlying skeleton we used a grid of rationally-placed surface markers, which were either distinct anatomical landmarks like the snout or were painted on the animal’s fur (Fig 1a, 14 landmarks and 29 spots total per animal, Supplementary Fig. 2). To individually tailor the skeleton to each animal we used gradient decent optimization to learn varying bone lengths and surface marker locations for each animal (see Methods “Learning bone lengths and surface marker positions”). Visible surface markers were manually annotated from a fraction of all recorded images to learn the skeleton that could then be used for all behavioral data acquired from that animal. As the rigid spatial relationship between surface markers and the underlying joints remained constant the algorithm could learn individual bone lengths as well as surface marker locations by adjusting both via gradient descent optimization (Supplementary Fig. 3). New poses were iteratively generated for each time point by applying a global translation to the generalized skeleton model and subsequently modifying positions of joints and rigidly attached surface markers by rotating each bone (Supplementary Video 1). Errors were established and minimized by projecting inferred 3D surface marker locations onto calibrated overhead camera sensors and subsequently comparing them to manually labeled ground-truth data (Supplementary Fig. 4).

To evaluate the accuracy of both the skeleton model and inference of joint positions over a large range of animal sizes, we obtained high-resolution MRI scans for each animal (Fig. 1f, N = 6 animals, see methods “Comparison of skeleton parameters with MRI data”) and aligned the skeleton model to measured positions of 3D surface markers (Fig. 1g). Errors for inferred spine and limb joint positions were low (Fig. 1h, 138 joint positions total, joint position error: 0.79 +/− 0.69 & 0.65 cm [mean +/− s.d. & median]) and inferred limb bone lengths and bone angles were not significantly different from those measured in MRI scans (Fig. 1i, 108 bone lengths total, range of measured bone lengths: 0.53 cm to 4.76 cm, bone length error: 0.46 +/− 0.34 & 0.36 cm [mean +/− s.d. & median], Spearman correlation coefficient: 0.75, two-tailed p-value testing non-correlation: 5.00×10^−21^; 84 bone angles total, range of measured bone angles: 4.13° to 123.77°, bone angle error: 27.80 +/− 18.98 & 26.72° [mean +/− s.d. & median], Spearman correlation coefficient: 0.47, two-tailed p-value testing non-correlation: 5.29×10^−6^). Together this demonstrated first, that the anatomically constrained skeleton model generated by our algorithm was highly accurate when compared with the animal’s actual skeleton across the range of animal sizes, and second, that accurate joint positions could be reconstructed in a single static pose from this approach.

### Accurate behavior reconstructions required both the temporal and anatomical constraints

To reconstruct behavioral sequences using the ACM, we first tracked 2D surface marker locations in the recorded movies using DeepLabCut^24^, which is specifically designed for surface landmark detection of laboratory animals. As the ACM contained both joint angle limits and temporal constraints, we evaluated the role of these by reconstructing poses without either the joint angle limits or the temporal constraints. The resulting temporal model, only temporally constrained, and the joint angle model, only constrained by joint angle limits, and the naïve skeleton model, constrained by neither, were compared to the ACM. To measure animal paw positions and orientations during gait we used a modified frustrated total internal reflection (FTIR) touch sensing approach^40,41^ (Fig. 2a-c, Supplementary Video 2) and compared these measurements to the paw positions and orientations inferred by each model (Fig. 2d,e, N = 6 animals, 29 sequences, 181.25 s per 145000 frames total from 4 cameras). The ACM produced significantly smaller positional errors compared to all other models (Fig. 2g, left; 10410 positions total; p-values of one-sided Kolmogorov-Smirnov test: ACM vs. joint angle model: 9.84×10^−21^; ACM vs temporal model: 4.38×10^−35^; ACM vs. naïve skeleton model: 9.03×10^−37^), whereas orientation errors were only significantly smaller when comparing the ACM to the temporal and naïve skeleton model (Fig. 2g, right; 7203 and 6969 orientations total for the ACM/anatomical model and the temporal/naïve skeleton model respectively; p-values of one-sided Kolmogorov-Smirnov test: ACM vs temporal model: 3.20×10^−39^; ACM vs. naïve skeleton model: 2.51×10^−50^). While orientation errors were significantly reduced by the anatomical constraints, including temporal constrains limited abrupt pose changes over time compared to either the naïve skeleton model or joint angle model (Fig. 2f, Supplementary Video 3-8). As a result, ACM-generated joint velocities and accelerations (Fig. 2h, 576288 velocities and accelerations total) were significantly smaller when compared to all other models (p-values of one-sided Kolmogorov-Smirnov test: ACM vs. joint angle model (velocity): numerically 0; ACM vs. temporal model (velocity): numerically 0; ACM vs. naïve skeleton model (velocity): numerically 0; ACM vs. joint angle model (acceleration): numerically 0; ACM vs. temporal model (acceleration): numerically 3.71×10^−90^; ACM vs. naïve skeleton model (acceleration): numerically 0). The temporal and anatomical constraints each had an advantage over the naïve skeleton model, and both constraints applied simultaneously improved positional accuracy as well as motion trajectories and prevented anatomically infeasible bone orientations and abrupt paw relocations. Moreover, the fraction of position errors exceeding 4 cm increased when constraints were not considered (fraction of errors exceeding 4 cm: ACM: 2.72%; joint angle model: 3.64%; temporal model: 4.42%; naïve skeleton model: 6.44%), and the same was observed for orientation errors exceeding 60° (fraction of errors exceeding 60°: ACM: 7.78%; joint angle model: 7.81%; temporal model: 17.77%; naïve skeleton model: 18.22%). Likewise, enforcing constraints also lowered the percentage of velocities exceeding 0.08 cm/ms (fraction of errors exceeding 0.08 cm/ms: ACM: 3.29%; joint angle model: 13.49%; temporal model: 3.28%; naïve skeleton model: 13.85%) and accelerations exceeding 0.02 cm/ms^2^ (fraction of errors exceeding 0.02 cm/ms^2^: ACM: 0.22%; joint angle model: 23.43%; temporal model: 0.25%; naïve skeleton model: 24.55%). To test ACM robustness to missing surface markers, position errors were calculated for inferred paw positions from data in which surface markers were undetected (Fig. 2i, 2797 position errors total). Compared to all other models the ACM produced significantly lower errors (p-values of one-sided Kolmogorov-Smirnov test: ACM vs. joint angle model: 9.67×10^−23^; ACM vs temporal model: 2.83×10^−22^; ACM vs. naïve skeleton model: 3.91×10^−47^) as well as the smallest number of error values above 4 cm (ACM: 9.36%; joint angle model: 11.61%; temporal model: 13.72%; naïve skeleton model: 19.12%). Paw placement errors increased the longer a surface marker remained undetected for the ACM and the naïve skeleton model (Fig. 2j, linear regression: ACM: slope: 1.49 cm/s, intercept: 1.13 cm; naïve skeleton model: slope: 2.77 cm/s, intercept: 1.39 cm) and errors were significantly lower when comparing both models (p-values of one-sided Mann-Whitney rank test: ACM vs. naïve skeleton model: 3.91×10^−47^).

**Fig. 2.**
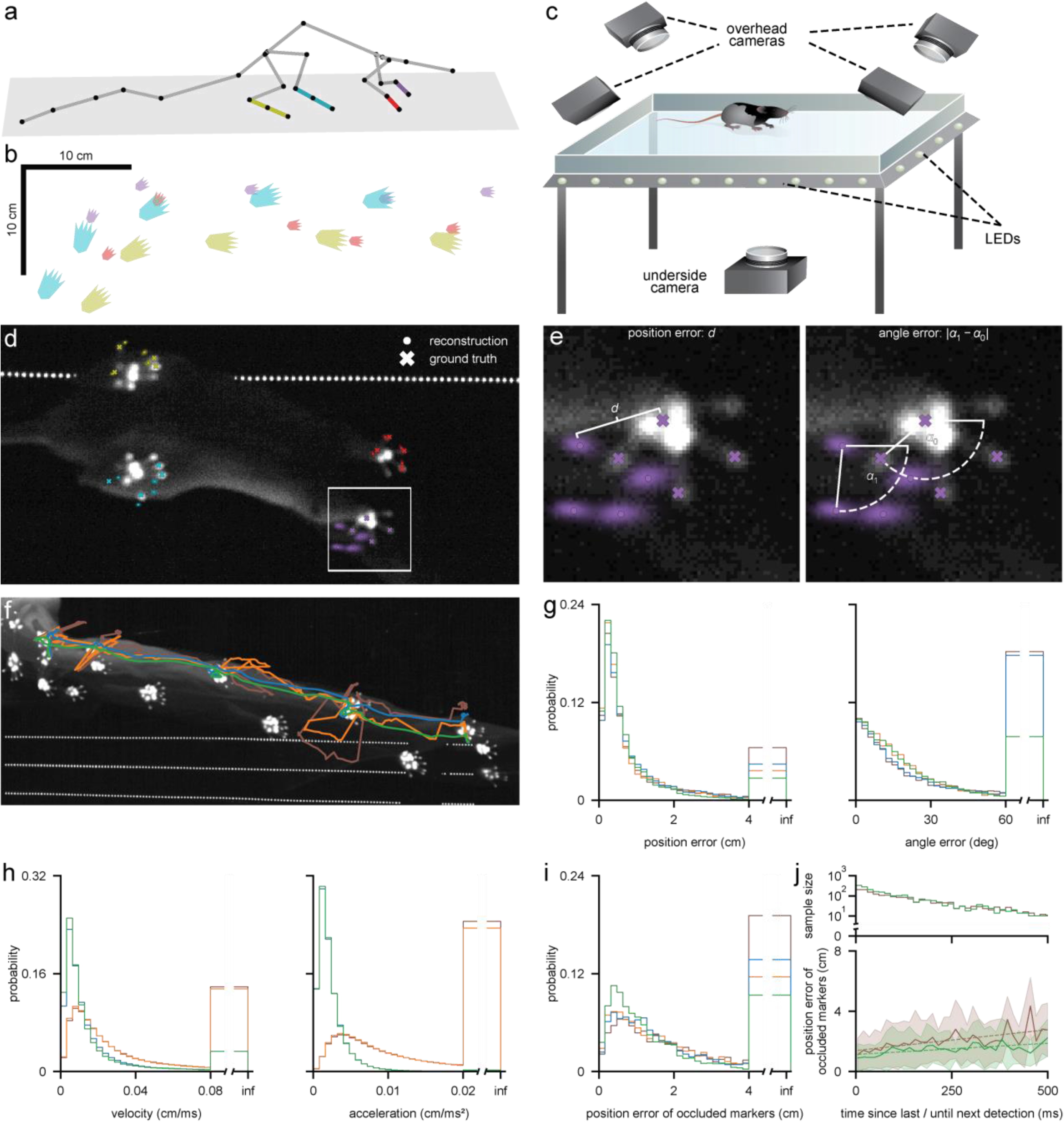
Comparison between inferred and measured paw positions during free behavior. **a**, Reconstructed animal pose based on a learned skeleton model with highlighted left front (purple), right front (red), left hind (cyan) and right hind paw (yellow). **b**, Reconstructed xy-positions of the paws during gait. Colors as in a. **c**, Schematic image of the FTIR touch sensing setup with one underneath and four overhead cameras. **d**, Single image from the underneath camera with reconstructed (x) and ground truth (filled circle) xy-positions of the paw’s centers and fingers/toes for the all four paws. Colors as in a. Large point clouds around landmark locations indicate high uncertainty. **e**, Enlarged view of the left front paw in d (white box) showing calculation of position error (left) and the angle error (right). **f**, Maximum intensity projection from the underneath camera of a 2.5 s long sequence with trajectories for the reconstructed xy-positions of the right hind paw using the ACM (green), temporal-(blue), joint angle-(orange) and naïve skeleton (brown) models. **g**, Probability histograms for paw position (left) and angle errors (right) comparing different model constraint regimes. Color-coding as in f. **h**, Probability histograms for paw velocities (left) and accelerations (right) comparing different model constraint regimes. Color-coding as in f. **i**, Probability histograms for paw position errors whereas only undetected surface markers are used for the calculation comparing different model constraint regimes. Color-coding as in f. **j**, Position errors of occluded markers (bottom) and corresponding binned sample sizes (top) as a function of time since last / until next marker detection comparing different model constraint regimes. Color-coding as in f. Sample sizes differ depending on whether reconstructed poses were obtained via the unscented RTS smoother (green) or not (brown).

### Kinetics of cyclic gait behavior

Smooth and periodic reconstruction of an animal’s average gait cycles during walking or running is only possible with robust and accurate tracking of animal limb positions. To establish whether the ACM could generate an average gait cycle from freely moving data, we next extracted individual gait cycles from multiple behavioral sequences (Fig. 3a,b, Supplementary Fig. 5, Supplementary Video 9-11) where joint velocities exceeded 25 cm/s (left; N = 2 animals, 27 sequences, 146.5 s per 58600 frames total from 4 cameras). The ACM extracted gait cycles were stereotypical and rhythmic (Fig. 3b,c), showing clear periodicity in autocorrelations of extracted limb movement (Fig 3c, left; damped sinusoid fit: frequency: 3.14 Hz, decay rate: 2.49 Hz, R^2^-value: 0.90) and a common peak for all limbs in Fourier transformed data (Fig. 3c, right; max. peak at 3.33 Hz, sampling rate: 0.83 Hz). Averaged ACM extracted gait cycles (Fig. 3d, left, Supplementary Fig. 6–9) were significantly less variable than those obtained from the naïve skeleton model (Fig. 3d, center) throughout the entire gait cycle (p-value of one-sided Mann-Whitney rank test: 1.40×10^−49^). When gait cycles were obtained from only tracking surface markers alone via a deep neural network without any form of underlying skeleton (surface model), high noise levels even made the periodic nature of the gait cycles vanish in its entirety (Fig. 3d, right). The robustness and accuracy of limb tracking was even more apparent when analyzing joint velocities (Fig. 3e-g), joint angles (Fig. 3h-j), and joint angular velocities (Fig. 3k-m), as traces generated without the ACM constraints were dominated by noise in individual examples (Fig. 3f,i,l, lower) and the cyclic nature of gait was less prominent when compared to traces obtained from the ACM (Fig. 3f,i,l, top). Consistent with this, ACM averaged traces (Fig. 3g,j,m, left, Supplementary Fig. 6–9) had significantly less variance compared to those obtained from the naïve skeleton model (Fig. 3g,j,m, right, Supplementary Fig. 6–9) for all metrics (p-values of one-sided Mann-Whitney rank test: velocity: 2.28×10^−55^; angle: 1.42×10^−55^; angular velocity: 1.44×10^−5^). Additionally, for all metrics the periodicity of the gait cycles in the form of equidistant peaks was more variable for the naïve skeleton model (12 peaks total; sampling rate: 10 ms; time difference between minimum/maximum peaks: position (min. peaks): 64.16 +/− 56.78 ms; velocity (max. peaks): 80.83 +/− 54.99 ms; angle (max. peaks): 74.16 +/− 33.53 ms; angular velocity (min. peaks): 53.33 +/− 47.78 ms [avg. +/− s.d.]), when compared to the ACM (12 peaks total; sampling rate: 10 ms; time difference between minimum/maximum peaks: position (min. peaks): 75.00 +/− 29.01 ms; velocity (max. peaks): 78.33 +/− 10.67 ms; angle (max. peaks): 78.33 +/− 23.74 ms; angular velocity (min. peaks): 75.00 +/− 10.40 ms [avg. +/− s.d.]). Together this shows that the ACM can objectively extract behaviors, such as gait, from freely moving animals and quantify complex relationships between limb-bones by inferring 3D joint positions over time as well as their first derivatives.

**Fig. 3.**
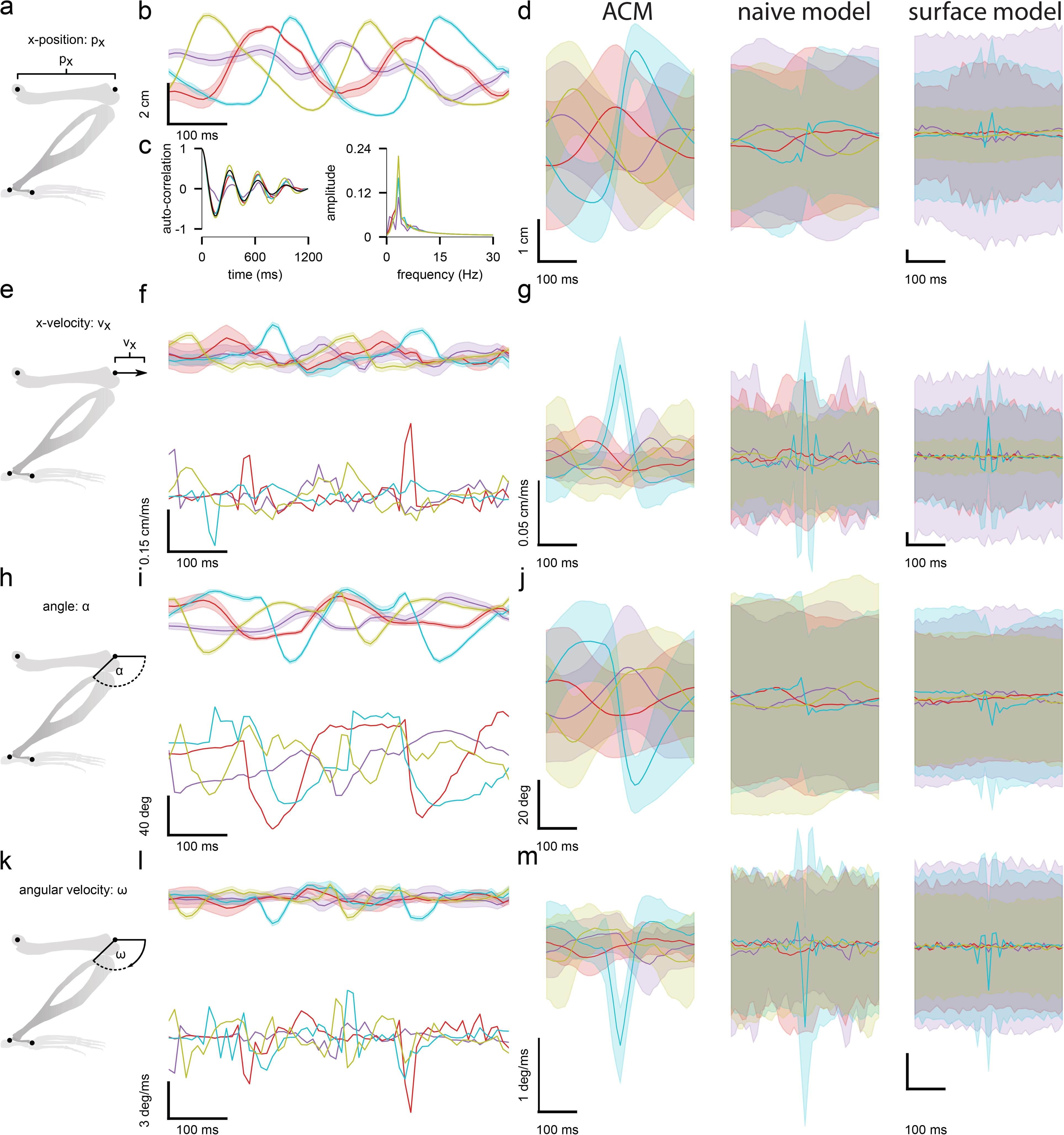
Influence of temporal and anatomical constraints on periodic gait cycles. **a**, Schematic of the normalized x-position for a single joint. **b**, Trajectories of the normalized x-position as a function of time for the left wrist (purple), right wrist (red), left ankle (cyan) and right ankle (yellow) joint during gait. **c**, Auto-correlations of the normalized x-position as a function of time (left) for four different limbs as well as a corresponding model fit via a damped sinusoid (black). Fourier transformed auto-correlations of all limbs (right) have their maximum peak at the same frequency. Colors as in b. **d**, Population averaged trajectories of the normalized x-position as a function of time for the ACM (left), the naïve skeleton model (center) and the surface model (right). Colors as in b. Trajectories of the ACM and the naïve skeleton model correspond to the 3D joint locations, whereas trajectories of the surface model correspond to the 3D locations of the associated surface markers. **e**, Schematic of the of the first temporal derivative of the normalized x-position (i.e. normalized x-velocity) for a single joint. **f**, Normalized x-velocity as a function of time for the ACM (top) and the naïve skeleton model (bottom) during gait (colors as in b). **g**, Population averaged trajectories of the normalized x-velocity as a function of time for the ACM (left), the naïve skeleton model (center) and the surface model (right). Colors as in b. Trajectories as in d. **h**, Schematic of the normalized bone angle for a single joint. **i**, Bone angle as a function of time for the ACM (top) and the naïve skeleton model (bottom) during gait (colors as in b). **j**, Population averaged trajectories of the bone angle as a function of time for the ACM (left), the naïve skeleton model (center) and the surface model (right). Colors as in b. Trajectories as in d. **k**, Schematic of the first temporal derivative of the bone angle (angular velocity) for a single joint. **l**, Angular velocity as a function of time for the ACM (top) and the naïve skeleton model (bottom) during gait (colors as in b). **m**, Population averaged trajectories of bone angular velocity as a function of time for the ACM (left), the naïve skeleton model (center) and the surface model (right). Colors as in b. Trajectories as in d.

### Kinetics of complex behavior

We next used the ACM to analyze motion kinetics and segment a more complex decision-making behavior, the gap crossing task, in which the distances between two separate platforms are changed forcing the animal to re-estimate the distance to jump for each trial (Fig. 4a). Reconstructed poses during gap-estimation and jump-behaviors consisted of sequences where animals either approached or waited at the edge of the track and jumped (N = 42, Supplementary Fig. 10, Supplementary Video 12,13) or reached with a front paw to the other side of the track before jumping (N = 2, Supplementary Video 14,15). As with the inference of paw placement during gait (Fig. 2b,f), hind-paw spatial position could be inferred throughout the jump and compared to skeletal parameters during the behavior, such as the angle of the thoracolumbar joint at jump onset compared to the paw positions upon landing (Fig. 4b; 44 trials, N = 2 animals). As rats jumped stereotypically, we next tested whether the jump-related pattern of movements could be analyzed using the ACM to objectively define decision points in the behavior, such as time of jump, from each individual trial. The changes in joint angles in the spine segments and hind limbs around the time of the jump were highly consistent. Averaging these joint angles to give an averaged joint-angle trace provided a metric with a global minimum (Fig. 4c) during the jump that was independent of whether the animal crossed the gap immediately, paused and waited at the track edge or reached across the gap (Supplementary Fig. 11). This approach enabled objective identification of jump start-, mid- and end-points, from each individual jump. Traces of joint angles averaged across joints and trials (Fig. 4d) and average ACM poses (Fig. 4e) illustrate the consistency of the pose changes through the jump. We next used this to quantify relationships between joints and changes in joint kinetics throughout a jump sequence. Auto-correlations for spatial and angular limb velocities allowed quantification of the interdependency of joint movements at any point within the jumping behavior, for example at the start-point of a jump (Fig. 4f,g). This displayed relationships like a significant correlation between the spatial velocity of the right elbow and wrist joints (Fig. 4h left, Spearman correlation coefficient: 0.95, two-tailed p-value testing non-correlation: 5.40×10^−24^), as well as joint interactions across the midline, such as a significant correlation between spatial velocity of the right and left knee joints (Fig 4h, right, Spearman correlation coefficient: 0.93, two-tailed p-value testing non-correlation: 6.79×10^−20^). As the animal jumped across the gap, changes in the bone angles and their derivatives (Fig. 4i) were correlated with distance that the animal jumped (Fig. 4j). For example, angular velocity of the thoracolumbar joint and vertical velocity (z-velocity) of the thoracocervical joint were significantly correlated with jump distance 205 ms and 175 ms respectively, before the animal landed (Fig. 4k,l, Spearman correlation coefficient: −0.73, two-tailed p-value testing non-correlation: 1.13×10^−8^, and 0.81, two-tailed p-value testing non-correlation: 1.12×10^−11^).

**Fig. 4.**
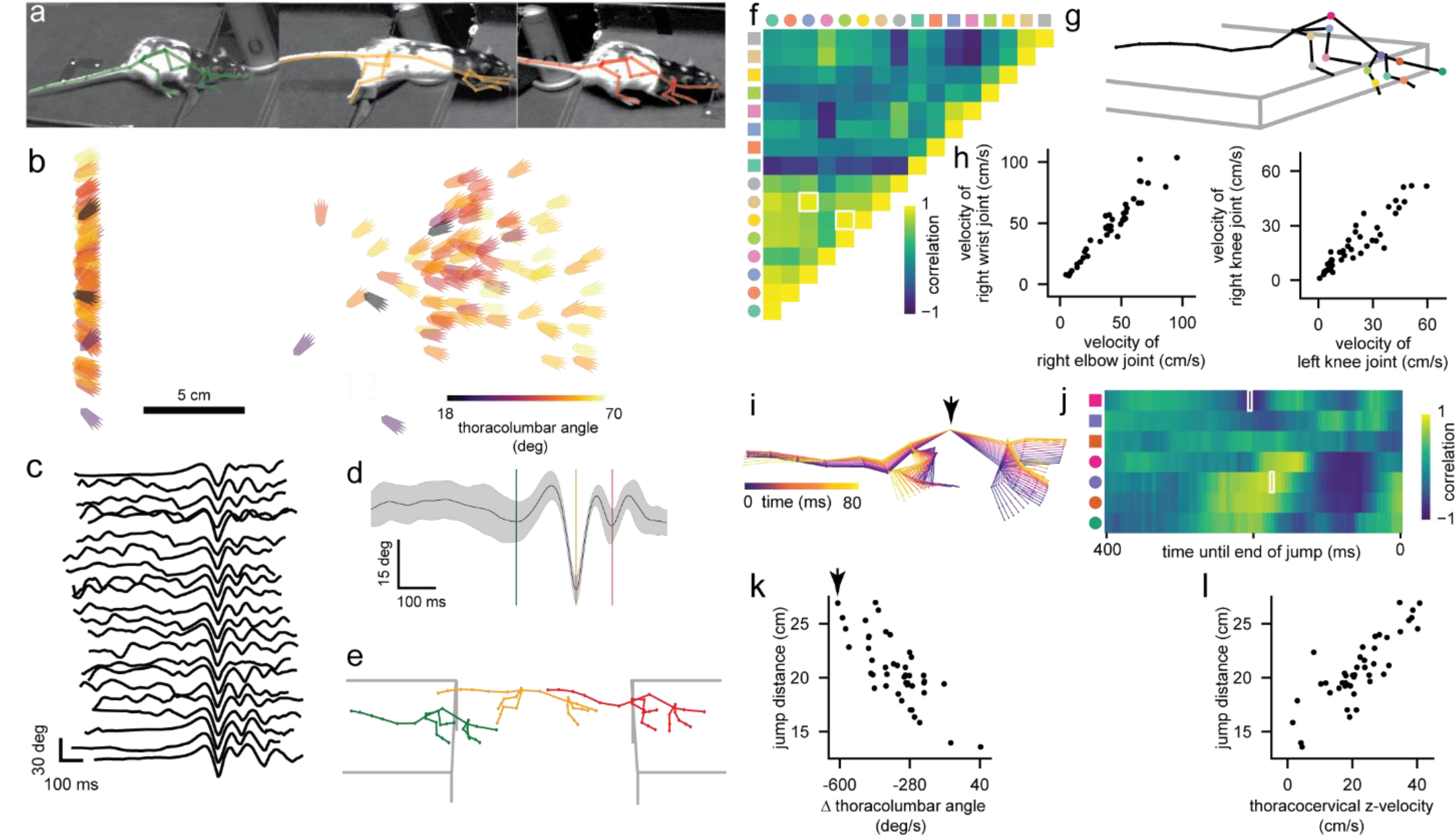
3D pose reconstruction of skeletons allows for detailed quantification of complex behavior. **a**, Images of a rat performing the gap crossing task for a trial. **b**, Reconstructed xy-positions of the hind paws at the start and end of the jump color-coded by the joint angle of the thoracolumbar joint for each gap crossing event of the population. **c**, Averaged joint-angle traces (spine and hind limb joint angles) from 22 out of 44 jump trials. **d**, Joint-angle trace averaged across joints and all jump trials. **e**, Average poses at the start-(green), mid-(orange) and end-point (red) of the jump from all jump trials. The three different time points are indicated by colored lines in d. **f**, Cross-correlation of the spatial and angular velocities of the limb joints at the start-point of a jump. Different marker shapes indicate whether rows/columns represent spatial or angular velocities (circles and squares respectively). Marker color corresponds to joint markers in g. **g**, Average pose at the start of a jump calculated from all jump trials. Joint colors are consistent with the marker colors in f and j. **h**, High correlation examples for spatial velocities of different limb joints as a function of each other for both animals. The data shown represents the correlation values highlighted in white in f. **i**, Overlaid poses of a single animal 240 ms to 160 ms before the end of a jump. Arrow indicates the thoracolumbar joint. **j**, Correlations of the z- and angular velocities of the head and spine joints for time points up to 400 ms before the end-point of a jump. Marker conventions as in f. **k**, Jump distance as a function of angular velocity of the thoracolumbar joint for both animals 205 ms before the end of the jump. Poses corresponding to the single data point highlighted with the arrow are shown in i. Displayed data represents the correlation value highlighted with a white rectangle in j. **l**, Jump distance as a function of z-velocity of the thoracocervical joint for both animals 175 ms before the end of the jump. Displayed data represents the correlation value highlighted with a white rectangle in j.

## Discussion

We developed an anatomically constrained model (ACM) for tracking skeletal poses of untethered freely-moving animals, at the resolution of single joints, that enabled the quantification of joint kinetics during gait and gap-crossing behaviors. From these kinetic measurements the ACM was able to build a comparative map of the kinetic-sequences throughout decision-making behaviors that could be compared to the behavioral outcome. Accurate generation of skeleton kinetics relied on incorporating skeleton anatomy, requiring smoothness of rotations and imposing motion restrictions of joints^27^, as animal poses are limited by both bone lengths and joint angle limits^26^. In addition, we generated ground truth data to quantify both the accuracy of the algorithm used to fit the model skeleton to the behavioral data and also the performance of the ACM at estimating limb and joint trajectories. Comparing the bone lengths of the fitted skeleton to the actual bone lengths measured from anatomical MRI scans for animals of a range of sizes we showed that accurate model fits could be obtained for animals with an order of magnitude difference in weight, with equally good fitting results independent of animal size. By directly measuring the animals paw positions and comparing with positions returned by the ACM, we showed that the combination of both anatomical and temporal constraints significantly reduced the errors relative to the naïve skeleton model or either constraint alone. This combination allowed accurate estimation not only of the location and orientation of the paws but also the accelerations and velocities of the joints during the measured behaviors. The ACM was capable of accurately quantifying limb kinetics during cyclic gait behaviors and more complex behaviors, such as gap-crossing, even when limbs were partially occluded.

Lastly, the ACM remained accurate over a large range of animal sizes, 72 g – 735 g, with the expectation that the ACM approach would also work for smaller rodents, such as mice. Our approach ushers in a suite of new possibilities for studying the biomechanics of motion during complex behaviors in freely-moving animals and complements recent developments in detailed surface tracking^42^. It opens up future investigations to also model forces applied by tendons and muscles^26,27^ and starts bridging the gap between neural computations in the brain^3–8,13^ and the mechanistic implementation of complex behavior^9–11^, such as rodent emotion^43^.

Recently, various studies relying on deep neural networks approached the problem of detecting an animal’s pose in the form of 2D features from an image without anatomically constrained skeleton models^21,23,24^. 3D poses can be inferred from these 2D features by means of classical calibrated camera setups^44^, however the 2D detection in one camera image does not benefit from the information from other cameras and the triangulation may suffer from resulting mislabeling of 2D features as well as missing detections due to occluded features. A recent approach^32^ overcomes many of these issues by mapping from recorded images directly to 3D feature locations, again using deep learning, and is capable of classifying animal behaviors across many species^32^. However, it does not possess explicit inherent temporal connections between frames and thus no persistent skeletal model with fixed bone lengths over time or anatomical constraints on joint angles. In contrast, the ACM uses a different approach: With DLC^24^ we used an existing method to detect 2D anatomical markers and inferred 3D positions and kinetics of movement with the RTS smoother based on anatomical constraints and mechanistic knowledge of bone rotations^26,27^, considering the trajectory of 3D positions over time. While the goal of the current study was to infer skeletal kinematics of freely behaving animals but not real-time behavior tracking^23,45^, we expect future work in the field of 3D animal pose estimation to combine both supervised learning techniques^32,42^ and mechanistic model constraints^26,27^, to simultaneously capitalize on their different strengths, e.g. by applying a smoother with anatomical knowledge like the ACM directly to 3D positions from an image-to-3D framework^32^. Our approach has the capacity to extend existing methods and not only to enhance the detail in which animal behavior can be studied and quantified, but it also provides an objective and accurate quantification of limb and joint positions for comparison with neuronal recordings.

## Acknowledgments

We thank David Greenberg for developing initial video acquisition software, Florian Franzen for help with an external camera trigger, Jan-Matthis Lückmann for initial code for reading single frames, Michael Bräuer, Rolf Honnef, Michael Straussfeld and Bernd Scheiding from the mechanical workshop for fabrication of the setup components, Kristina Barragan, Abhilash Cheekoti, Nada Eiadeh, Gizem Görünmez, Yolanda Mabuto, Anastasiia Nychyporchuk, Aarya Pawar and Nurit Zorn for manually labelling of images and Julia Kuhl for illustrations. Funding was obtained from Stiftung caesar and the Max Planck Society. A.M. is a graduate student with the International Max Planck Research School (IMPRS) for Brain and Behavior. K.S. and E.L. were funded in part by DFG Reinhard Koselleck Project, DFG SCHE 658/12. J.H.M. was supported by the German Research Foundation (DFG) through Germany’s Excellence Strategy (EXC-Number 2064/1, Project number 390727645).

## Author contributions

Development of anatomically constrained model concept: A.M., J.H.M., and J.N.D.K. Algorithm design and implementation: A.M. and J.H.M. Experimental design and setup: A.M., K-M.V., D.J.W., J.S., and J.N.D.K. Animal preparation and data collection: A.M. and D.J.W. MRI sequences and data collection: A.M. E.L. and K.S. Analysis design and implementation: A.M. and J.N.D.K. Manuscript preparation: A.M. and J.N.D.K.

## Methods

### Obtaining video data of behaving animals

All experiments were performed in accordance with German guidelines for animal experiments and were approved by the Landesamt für Natur, Umwelt und Verbraucherschutz, North Rhine-Westphalia, Germany. Six Lister Hooded rats (Charles River Laboratories), weighting 174 g (animal #1), 178 g (animal #2), 71 g (animal #3), 72 g (animal #4), 735 g (animal #5) and 699 g (animal #6) on the day of the experiment, were used. Anatomical landmarks for tracking limb and body positions consisted of black or white ink spots (5-8 mm diameter, black markers: Edding 3300, white markers: Edding 751, Edding, Ahrensburg, Germany) which were painted onto the fur in a stereotypical pattern that was near-symmetrical around the animals’ mid-sagittal axis (Supplementary Fig. 2). For application of the anatomical markers, animals were anesthetized with isoflurane (2-3%) and body temperature maintained around 37.5°C using a heating pad and temperature probe. After this labeling procedure the animals were allowed to recover for approx. 45 min before datasets were acquired on a gap-crossing track and open arena. The open arena was 80×105 cm^2^ with 50 cm high walls colored gray to promote contrast with the animals and markers. The gap-crossing track consisted of two 50×20 cm^2^ platforms, mounted 120 cm off the ground on a slide mechanism to allow manual adjustment of the distance between the platforms in the range from 0 to 60 cm. The platforms were positioned such that with the gap closed they met along one of the 20 cm edges. The edges of the platform, apart from the edge along which the two platforms met, were equipped with a 2.5 cm tall wall. The floor of the platforms was covered with a layer of neoprene material to promote a secure grip for the animals’ feet. A water delivery spout was located in the center of the 20 cm track edge opposite of where the platforms met. To encourage gap-crossing behavior, animals were water-restricted, having full access to water two days per week, and otherwise having access to water only on the gap-crossing track. Fifty to one hundred microliters of water was available at the delivery spout after each successful cross of the gap. Animals received a minimum of 50% of their daily *ad libertum* water consumption either during the training or recording sessions or as a supplement after the last session of the day if they had not already consumed at least this amount. Gap-crossing training commenced approximately two weeks prior to the recording, and consisted of two daily sessions. Gap distances were pseudo-random, with the gap distance reduced in cases where the animal refused to cross. Both setups were homogeneously illuminated using eight 125 cm long white LED strips with 700 lm/m (PowerLED, Berkshire, United Kingdom), arranged equidistantly in a patch of 125×80 cm^2^ and 125×55 cm^2^ at a distance of 130 cm and 150 cm above the ground of the open arena and the gap-crossing track, and data were acquired using four synchronously triggered digital cameras (ace acA1300-200um, Basler, Ahrensburg, Germany) mounted above the setups and set in such a way that all parts of the setup were covered by at least two cameras, with the majority of both setups covered by all four. Datasets consisted of 1280×1024 px^2^ image frames with an acquisition time of 2.5 ms recorded at 100 Hz for the gait dataset in the open arena and 200 Hz for the gap-crossing dataset. For quantification of the animals’ foot positions we used a custom-made FTIR plate, consisting of a single sheet of 60×60 cm^2^, along the edges of which an IR-LED strip (Solarox 850 nm LED strip infrared 850 nm, Winger Electronics GmbH & Co. KG, Germany) was mounted such that IR light could propagate through the FTIR plate from two opposing sites. Animal position data were acquired from the overhead cameras at 200 Hz for these experiments, and paw placements were recorded using two additional cameras (ace acA1300-200um, Basler, Ahrensburg, Germany), synchronized with the overhead cameras, mounted underneath the plate and equipped with infrared-highpass filters (Near-IR Bandpass Filter, part: BP850, useful range: 820-910 nm, FWHM: 160 nm, Midwest Optical Systems, Inc., Palatine, USA). These cameras were set to acquire 1280×1024 px^2^ frames with an acquisition time of 2.5 ms recorded at 200 Hz. The total FTIR dataset consisted of 29 sequences with a total of 36250 frames in each of the four cameras and a total duration of 181.25 s. The gait dataset consisted of 27 sequences with a total of 14650 frames in each of the four cameras and a total duration of 146.5 s. The gap-crossing dataset consisted of 44 sequences with a total of 8800 frames in each of the four cameras and a total duration of 44 s.

### Obtaining MRI scans to evaluate learned skeleton models

To locate labeled surface markers, custom-made MRI markers (Premium sanitary silicone DSSA, fischerwerke GmbH & Co. KG, Waldachtal, Germany) were attached to the respective positions on the surface of the animals’ bodies. MR imaging was performed at a field strength of 3T (Magnetom Prisma, Siemens Healthineers, Erlangen, Germany), using the integrated 32-channel spine coil of the manufacturer. The data was acquired *ex vivo* in six rats using a 3D turbo-spin-echo sequence with variable-flip-angle echo trains (3D *TSE*-*VFL*). Detailed MR protocol parameters for 3D TSE-VFL imaging with a turbo factor of 98 were as follows: a repetition time of 3200 ms, an effective echo time of 284 ms, an echo train duration of 585 ms, and an echo spacing of 6.3 ms using a readout bandwidth of 300 Hz/px for one slab with 208 slices covering the whole rat at an isotropic resolution of 0.4×0.4×0.4 mm^3^.

### Calibrating multi-camera setups

We based the calibration of multiple cameras on a pinhole camera model with 2nd order radial distortions (Supplementary Text) and OpenCV^46^ functions for detection of checkerboard corners. The checkerboards we used had additional ArUco^47^ markers printed on them. To obtain the calibration an objective function penalizing mismatches between detected and projected corners was defined and minimized via gradient descent optimization using the Trust Region Reflective algorithm^48^ (Supplementary Text).

### Defining a 3D skeleton model

The generalized skeleton model consisted of joints, modeled as vertices, and inter-joint segments, modeled as edges and which could represent multiple bones from the true skeleton (Supplementary Fig. 1). The front limbs were modeled as four edges, representing the clavicle, humerus, radius/ulna and metacarpal/phalanges. The associated vertices corresponded to the shoulder, elbow and wrist, with the last vertex representing the tip of the middle phalanx. The hind limbs were modeled as five edges representing the pelvis, femur, tibia/fibula, tarsus and phalanges, with the associated vertices representing the hip, knee, ankle and metatarsophalangeal joints, with the last vertex representing the tip of the middle tarsal. The tail was modeled as five edges and five vertices, with the last vertex representing the tip of the tail. The spine was modeled as four edges, representing the cervical, thoracic and lumbar spinal regions and the sacrum, with three intervening vertices. The head was modeled as a single edge, with a vertex at the tip to represent the nose, and a second vertex representing the joint to the first cervical vertebra. The resting pose of the 3D skeleton model was defined as the pose generated by the pose reconstruction scheme, when all the parameters encoding bone rotations were set to zero. In this pose all edges (i.e. bones) pointed towards the positive z-direction of the right-handed world coordinate system, except the four edges approximating the clavicle/collarbone and sacrum/pelvis, where edges of the right limb faced towards and edges of the left limb faced against the positive x-direction of the world coordinate system (Supplementary Fig. 1). The configuration of these four edges was also kept constant during pose reconstruction, so that edges representing the cervical and lumbar vertebrae were always orthogonal to the edges representing the clavicle and sacrum. The y-coordinates of all vertices were equal to zero, locating the entire 3D skeleton model in the world’s xz-plane while situated in the resting pose. Besides the world coordinate system each edge also had its own right-handed coordinate system located at the start vertex of the corresponding edge, e.g. the coordinate system of the edge representing the left humerus was located at the position of the vertex representing the left shoulder joint. The z-direction of these edge coordinate systems were always identical to the direction in which the associated edges faced. Additionally, anatomical rotations were defined in the edge coordinate systems, so that a rotation around the x-direction became equivalent to flexion/extension, rotations around the y-direction were identical to abduction/adduction and a rotation around the z-direction coincided with internal/external rotation.

### Constraining poses based on joint angle limits

We implemented joint angle limits based on measured minimum and maximum values for flexion/extension, abduction/adduction and internal/external rotation in domestic house cats^21^. A comparable set of measured values is to our knowledge not available for rat. For vertices approximating head, spine or tail joints data for joint angle limits was not available, so that we modeled corresponding edges without the capacity to rotate around the z-direction of their associated edge coordinate systems, whereas joint angle limits for rotations around the x- and y-direction were set to +/− 90°. This allowed a respective child-vertex to reach any point within an area spanned by a hemisphere pointing in the direction of the associated edge with radius identical to the length of this edge. Since the resting pose of our 3D skeleton model was not necessarily identical to the pose in which the published joint angle limits were measured in, we calculated the correct rotational limits which were consistent with our resting pose based on the mean of the published values. The resulting joint angle limits were set as follows:

| joint | x (°) | y (°) | z (°) |
| --- | --- | --- | --- |
| left shoulder | [25,205] | [-85,25] | [-35,35] |
| right shoulder | [25,205] | [-25,85] | [-35,35] |
| left elbow | [2.5,145] | [0,0] | [-100,45] |
| right elbow | [2.5,145] | [0,0] | [-45,100] |
| left wrist | [-135,35] | [-12.5,37.5] | [0,0] |
| right wrist | [-135,35] | [-37.5,12.5] | [0,0] |
| left hip | [35,195] | [-65,25] | [-85,40] |
| right hip | [35,195] | [25,65] | [-40,85] |
| left knee | [-145,15] | [0,0] | [0,0] |
| right knee | [-145,15] | [0,0] | [0,0] |
| left ankle | [-10,145] | [0,0] | [0,0] |
| right ankle | [-10,145] | [0,0] | [0,0] |
| left metatarsophalangeal | [0,0] | [0,0] | [-15,35] |
| right metatarsophalangeal | [0,0] | [0,0] | [-35,15] |

While the published joint angles referred to Euler angles, we used Rodrigues vectors to parameterize rotations (Supplementary Text), since the latter are better suited for pose reconstruction^49^. However, both parameterizations become identical when only a single type of rotation, e.g. flexion/extension, is present at a vertex, which was the case for the measured joint angles^49^. Parameterizing rotations with Rodrigues vectors therefore allowed us to obtain smooth transitions between different types of bone rotations.

### Constraining surface marker positions based on body symmetry

When learning surface maker positions and bone lengths we constrained the former to comply with the symmetrically applied surface marker pattern by enforcing box constraints for each spatial dimension, e.g. markers on the left side of an animal were prevented from being placed on the right side. This reduced the total number of free parameters during learning. To reduce this number further we also mirrored surface marker positions in the yz-plane of the associated edge coordinate system when there was a left/right correspondence, i.e. we only learned surface marker positions for the left side which then also determined right-sided surface maker positions due to the mirroring (Supplementary Text). Resulting box constraints for central and left-sided surface maker locations were then defined in the coordinate system of the associated edges and set as follows (Supplementary Fig. 1,2):

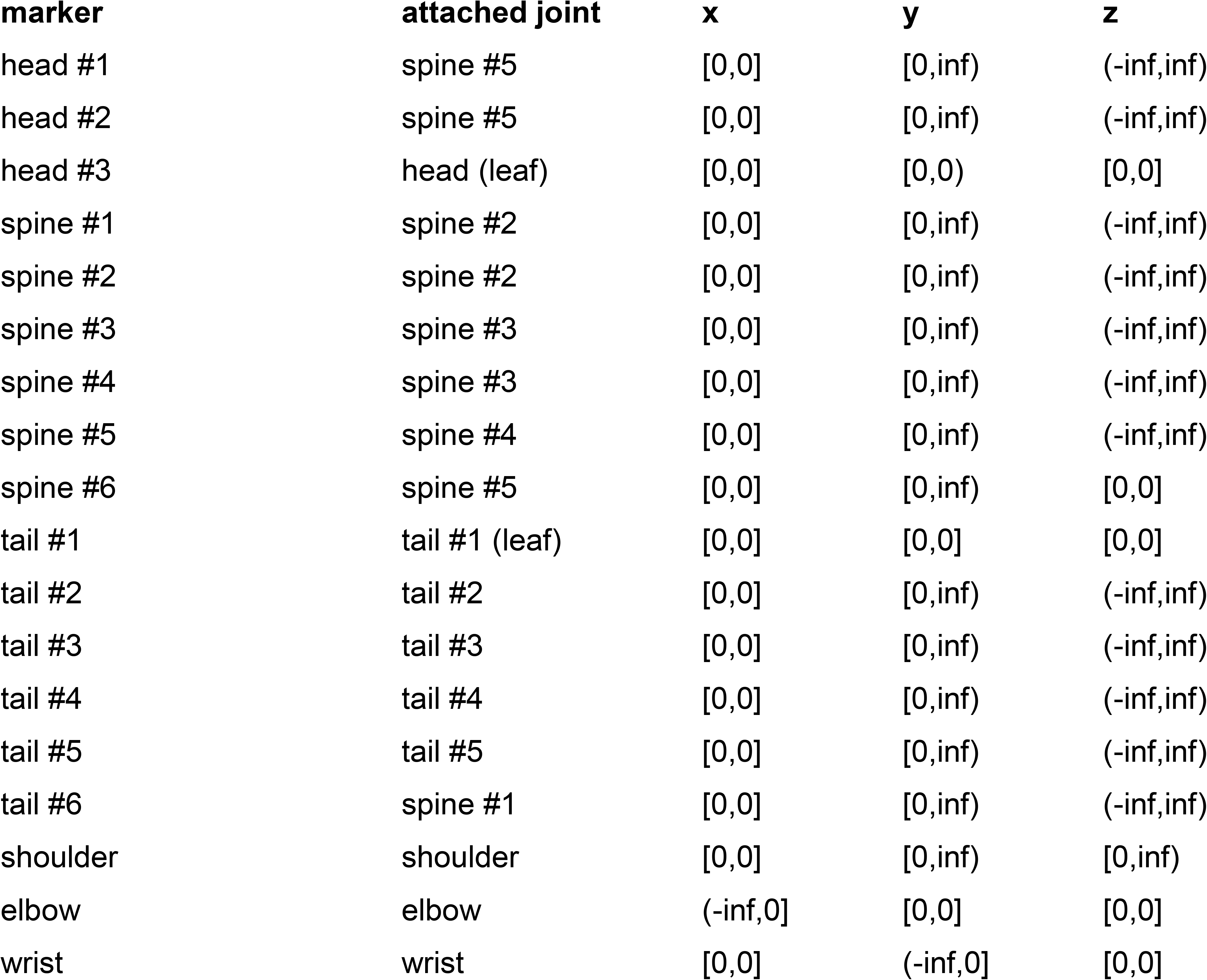
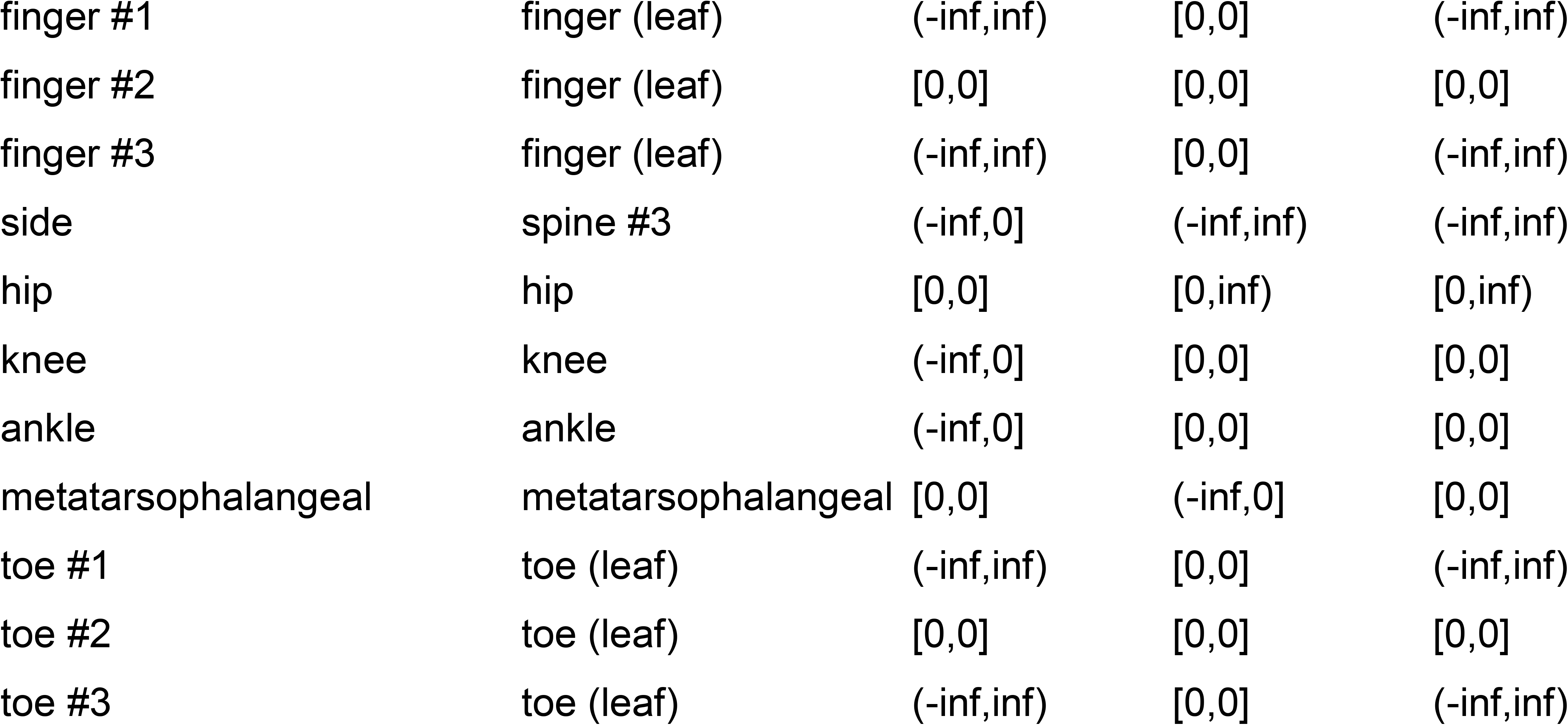

The only exception from this was the upper bound of the left-sided surface marker on the shoulder in z-direction for the two large animals (animals #5 and #6), which was also set to 0 in order to prevent the bone lengths of the collarbones to become zero during learning.

### Constraining bone lengths based on allometry

We applied loose constraints on the length of limb bones based on the published linear relationships between body weight and bone lengths in rats^35^. The lengths of the following list of limb bones were constrained according to measured slope estimates^35^. Box constraints for bone lengths were calculated from weight-matched lengths plus or minus 10 times the standard deviation, based on the following proportionality factors:

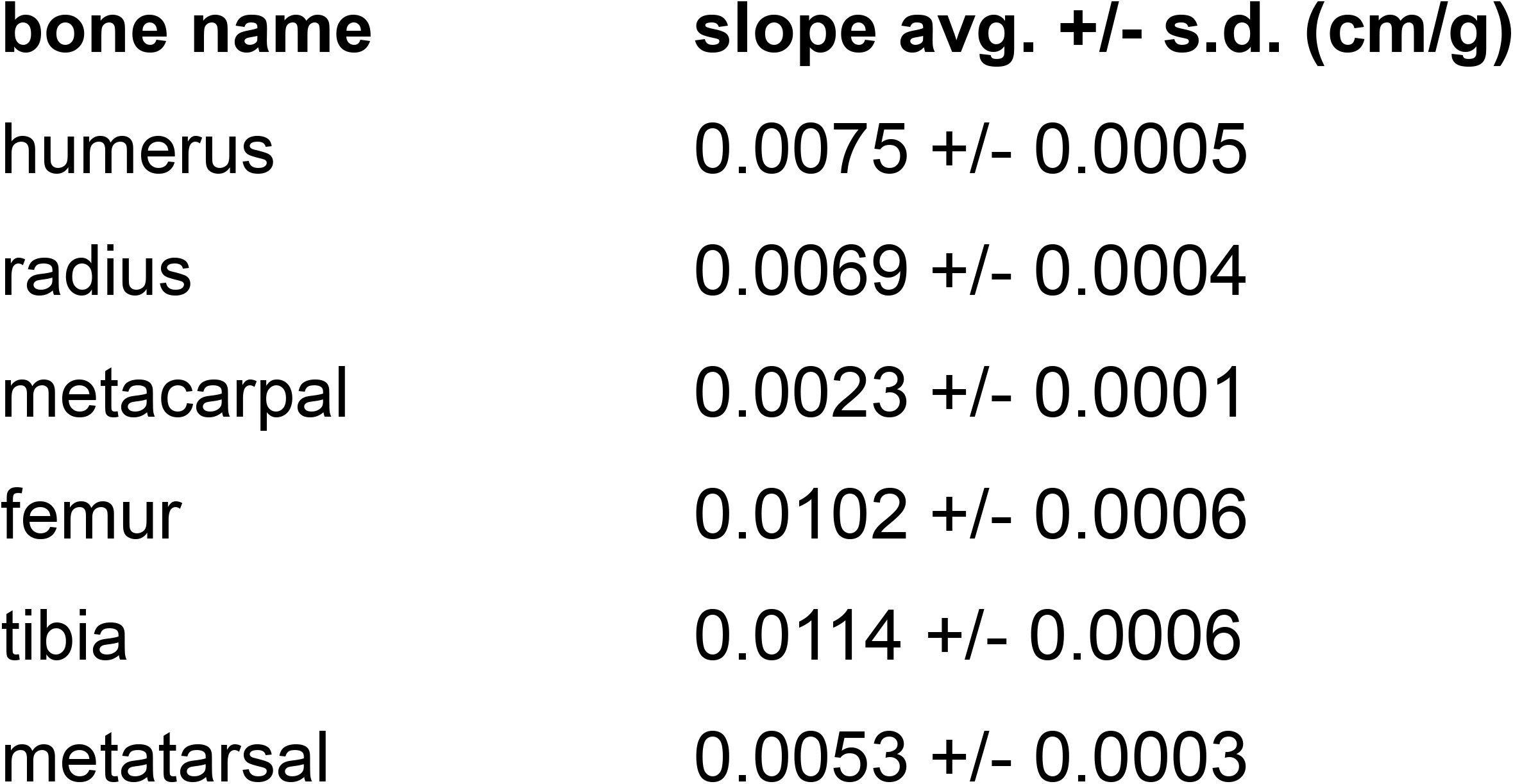

For bones that were not part of the limbs no constraints were enforced, such that corresponding box constraints were set to [0,inf). To ensure that bone lengths of the left and right limbs were identical, we only learned bone lengths of the left-sided limbs, which then also determined right-sided limb bone lengths (Supplementary Text).

### Training deep neural networks to detect 2D locations of surface markers

To automatically detect 2D locations of surface makers we used DeepLabCut^24^. For each rat in each dataset an individual neural network was trained on manually labeled images obtained from four different cameras, six trained networks in total. For each image that was used for training a background-subtracted image was generated by subtracting the image acquired 200 ms prior to the frame of interest for the FTIR and gap-crossing datasets and 125 ms prior for the gait dataset. Subsequently, approximate 2D locations of the recorded rats on the background-subtracted images were obtained by calculating the median indices of pixels above a threshold-value of 5 times the standard deviation of each pixel, where the standard deviations were calculated from the first 100 images of each recorded video, which were acquired with the arena or track empty and free of any moving objects. These 2D locations where then used as a center-point to crop the original images to 600×600 px^2^. To minimize the influence of visible movements of the experimentors on this center-point detection in the recorded FTIR data set, pixel values of pixels, which did not show the FTIR plate, were set to zero for the recordings of animals #3 to #6. For the FTIR datasets the networks were trained on 4068 images for animal #1, 3980 images for animal #2, 752 images for animal #3, 1100 images for animal #4, 992 images for animal #5 and 1128 images for animal #6. For the gait and gap-crossing datasets 2404 and 3608 images were used respectively for each analyzed animal (animal #1 and #2). Resulting images that did not contain any manually annotated 2D positions of surface markers due to the preprocessing steps not leading to correct cropping, were not used during training. We used DeepLabCut’s default settings, with the only two exceptions being that we changed the network architectures to ResNet-152 and enabled mirroring of images for which we paired surface markers with a left/right correspondence^50^. Training was conducted via DeepLabCut 2.1.6.4 downloaded from GitHub (https://github.com/DeepLabCut/DeepLabCut/commit/2f5d32884da2e5c3e4b6ef2a2126f6bb61579060). Once the networks were trained, we used them to obtain 2D locations of surface markers for images of analyzed behavioral sequences, where we set DeepLabCut’s pcutoff-parameter^50^ to 0.9 and treated detected marker positions below this value as missing measurements.

### Performing probabilistic pose reconstruction

To perform probabilistic 3D pose reconstruction, which allows for generating poses using non-linear mathematical operations and where information of an entire behavioral sequence is processed, we implemented an unscented Rauch-Tung-Striebel (RTS) smoother^36,37^, whose fundamental principles are based on the ordinary Kalman filter formulation^38^. In this approach, time series data is modelled as a stochastic process generated by a state space model where at each time point hidden states give rise to observable measurements and fulfill the Markov property, i.e. each hidden state only depends on the preceding one (Supplementary Fig. 12). This formalism allowed us to represent each pose as a low-dimensional state variable, corresponding to the location of a reconstructed skeleton as well as the individual bone rotations (dimension of hidden state variable: 50; 3 variables for 3D location of the skeleton plus 47 variables for bone rotations). The measurable 2D locations of surface markers (which were given by the outputs of the trained neural network) had a higher dimensionality and were represented via measurement variables (dimension of measurement variable: maximal 344; 43 surface markers times 4 cameras times 2 variables for the 2D location of a surface marker). We assumed the hidden states to be (conditionally) normally distributed, whereby temporal constraints are implicitly modeled through the transition kernel of the Markov process (i.e. the probabilistic mapping between one state and the next). Our formalism allows for non-linearities in our pose reconstruction scheme, e.g. introduced by the usage of trigonometric functions when applying bone rotations. The unscented RTS smoother can be used to perform probabilistic pose estimation in such a nonlinear state space model, considering both past and future (Supplementary Text). We learned the unknown model parameters (i.e. the initial mean and covariance of the state variables as well as the covariances of the transition and measurement noise) via an expectation-maximization (EM) algorithm^39^ (maximal 2944 model parameters total; 50 parameters for mean of initial hidden state variable plus 1275 parameters for covariance matrix of initial hidden state variable plus 1275 parameters for covariance matrix of transition noise plus maximal 334 parameters for diagonal covariance matrix of measurement noise), which aims to maximize a lower bound of the state space model’s evidence, i.e. the evidence lower bound (ELBO), accounting for each pose within a behavioral sequence (Supplementary Text). This is achieved by alternating between an expectation step (Supplementary Text), in which we obtain the expected values of the state variables given a fixed set of model parameters via the unscented RTS smoother, and a maximization step (Supplementary Text), in which these model parameters are updated in closed form in order to maximize the ELBO^51^. After convergence of the EM algorithm, final poses were obtained by applying the unscented RTS smoother using the learned model parameters.

### Accounting for missing measurements during pose reconstruction

Detecting 2D positions of surface markers via a trained deep neural network was not always successful, e.g. due to marker occlusions. As a result, we only had access to different subsets of all 2D positions during smoothing. This forced us to apply modifications to the plain unscented RTS smoother formulation and the EM algorithm, i.e. we set rows and/or columns of the measurement covariance matrices to zero during the filtering path of the smoother^52,53^ and proceeded equivalently with the covariance matrix of the measurement noise when maximizing the ELBO (Supplementary Text).

### Enforcing joint angle limits during pose reconstruction

The plain formulation of the unscented RTS smoother does not account for box constraints, so that state variables representing bone rotations are not bounded. To still allow for anatomically constrained pose estimation we instead optimized unbound state variables, which could be mapped onto the correct lower and upper bounds for joint angle limits via sigmoidal functions, i.e. error functions (Supplementary Text). These functions had slope one at the origin and were asymptotically converging towards the lower and upper bounds of the respective joint angle limits.

### Learning bone lengths and surface marker positions

To learn bone lengths and surface marker positions we simultaneously fitted our generalized 3D skeleton model to manually labeled 2D positions of surface markers at different time points for each animal. Fitting of the generalized 3D skeleton model was achieved via gradient decent optimization using the L-BFGS-B algorithm^54^ in order to minimize an objective function, which penalized mismatches between manually labeled 2D locations of surface markers and those generated via the pose reconstruction scheme (Supplementary Text). Bone lengths, surface marker positions and pose parameters were optimized, while only the pose parameters were unique for every time point and the rest were shared throughout the entire sequence. For this we used sequences of freely-behaving animals recorded via four different cameras totaling to 2404 training frames for animal #1 and #2, 752 training frames for animal #3, 1100 training frames for animal #4, 992 training frames for animal #5 and 1128 training frames for animal #6. Bone lengths were initialized by the mean of their upper and lower bounds or zero when there were no constraints and surface marker positions were initialized to be identical to the joints they were attached too. Initial poses were identical to the resting pose but global skeleton locations and rotations were adjusted prior to the fitting to loosely align with the locations of an animal’s body as seen by the cameras. Once values for bone lengths and surface marker positions were learned, we used them for all further pose reconstructions.

### Comparison of skeleton parameters with MRI data

To estimate the quality of these skeleton parameters, we aligned learned 3D skeleton models to manually labeled 3D locations of surface markers obtained from an MRI scan for each animal (Fig. 1b, bottom). To determine the 3D positions of the respective spine joints in the MRI scan, we counted vertebrae such that each modeled spine segment matched its anatomical counterpart with respect to the number of contained vertebrae^33^. One MRI surface marker was not recoverable in the MRI dataset from one animal (right metatarsophalangeal marker, animal #1), and in this case we labeled the 3D location on the animal’s body closest to the position of the missing marker. Again, we used gradient decent optimization of an objective function, so that manually labeled 3D markers locations were matched with the ones given by our model. Skeleton parameters were kept constant and only pose parameters changed during optimization. All ground truth joint positions except those for the metatarsophalangeal joints could be identified manually in the MRI scan (4 joint locations total). These missing locations were assumed to be identical to the positions of the corresponding metatarsophalangeal markers.

### Defining four different models to evaluate the influence of anatomical and temporal constraints

In the ACM anatomical and temporal constraints were enforced and poses were reconstructed using the unscented RTS smoother together with the EM algorithm. This was also the case for the temporal model but joint angle limits of limb joints, which were not equal to [0°,0°], were set to [−180°,180°], effectively allowing full 360° rotations at the respective joints. Pose parameters for these two models were initialized by fitting the pose of the first time point of a behavioral sequence equivalently to how we learned the skeleton parameters with the only exception that automatically detected instead of manually labeled 2D locations of surface markers were used. The covariance matrices for the initial state variables as well as the state and the measurement noise, which were learned via the EM algorithm, were initialized by setting all diagonal entries to 0.001 and off-diagonal entries to zero, while the latter were also kept constant for the measurement noise covariance matrix during the maximization step of the EM algorithm. For the joint angle and the naïve skeleton model, where only anatomical or no constrains were enforced, we did not use the unscented RTS smoother but reconstructed every pose in the same way we initialized the ACM and the temporal model. Here poses within a behavioral sequence at a certain time point were initialized with the reconstructed pose of the previous time point and joint angle limits of limb joints were set to [−180°,180°] in the naïve skeleton model.

### Evaluating pose reconstruction accuracy via a FTIR touch sensing system

To obtain ground truth data paw centers and three individual fingers/toes were manually labeled for each limb in every 40th frame of the FTIR dataset. Images from the calibrated underneath cameras were used and paw centers were identified as the interpolated intersection of the three fingers/toes. Manually labeled marker locations were then projected onto the surface of the transparent floor and xy-positions were calculated as the intersection between this surface and the corresponding epipolar lines. Paw positions and orientation errors where then calculated in the coordinate system of the transparent floor based on these xy-coordinates. Velocity and acceleration values for the four different models were derived from central finite differences (order of accuracy: 8) based on the reconstructed 3D positions of the metatarsophalangeal/wrist and finger/toe markers. Paw position errors of undetected markers where obtained by only using paw position errors of surface markers that were not detected by the trained neural network (i.e. pcutoff < 0.9). When ordering these errors according to the time spans that passed since a respective maker was successfully detected, differentiating between the ACM/temporal and joint angle/naïve skeleton model was necessary. As the unscented RTS smoother incorporates information from the past as well as from the future, time spans until the next detection need to be treated equally to time spans since the last detection, i.e. the direction of the time axis becomes irrelevant. For the ACM/temporal model time spans were calculated as the minimum of the time spans since the last or until the next detection, whereas for the joint angle/naïve skeleton model only time spans since the last detection were relevant, as the smoother was not used here. For the resulting analysis we only included errors for which the corresponding sample size was at least 10.

### Analyzing gait data

In order to extract gait periodicity, we normalized reconstructed poses by applying a coordinate transformation on 3D joint locations, such that the new origin was identical to the joint which connects lumbar vertebrae with the sacrum and the new x-direction pointed towards the xy-position of the joint linking cervical with thoracic vertebrae. Given the new joint coordinates we calculated normalized x-positions and bone angles as well as their first temporal derivatives (i.e. normalized x-velocity and angular velocity) of limb joints, where bone angles were defined as the angle between the new x-direction and a respective bone. To model auto-correlations of normalized x-positions, we minimized an objective function penalizing mismatches between data from the four different traces of each limb and the corresponding estimate calculated using a single damped sinusoid via gradient decent optimization. To obtain population averages of normalized x-positions, bone angles and their temporal derivatives, we detected midpoints of swing phases by identifying maximum peaks of normalized x-velocities above 25 cm/s. An individual trace was extracted containing data up to +/− 200 ms around each peak. These traces were then aligned with respect to their associated velocity peaks and then averaged across the entire population. To obtain traces from only tracking surface markers alone, 3D positions of surface markers where interpolated based on their inferred 2D locations given by the trained neural network, whereas for each 3D position only the two most likely 2D locations where taken into account, i.e. the two 2D locations with the highest pcutoff-value.

### Analyzing gap-crossing data

Each of the 44 gap-crossing sequences was 1 s long and contained 200 frames per camera totaling 35200 frames. Due to the limited number of gap-crossing events and recorded frames, we used 20% of the frames to train the neural network, i.e. we took every 5th frame of the recorded gap crossing sequences for its training and deployed it to automatically process all frames once training was completed. Similar to the analysis of gait data, velocity values were derived from central finite differences (order of accuracy: 8) of reconstructed 3D joint positions and joint angles were defined as the angle between two connected bones. To obtain start-, mid- and end-point for each jump we averaged joint angles of all spine and hind limb joints and identified the time point where this averaged metric reached its global minimum for each gap-crossing sequence. The averaged metric was characteristic for each jump, i.e. distinct peaks were always present in the following order: local minimum, local maximum, global minimum, local maximum, local minimum. This allowed us to extract the start- and end-point of each jump by finding the first and last local minimum of this sequential pattern. Resulting jump start- and end-points were in close agreement with those obtained from manual assessments of gap crossing sequences by a human expert. Jump distances were calculated as the absolute xy-difference of the average hind paw positions, i.e. average of left/right ankle, metatarsophalangeal and toe joint positions, at the start- and end-point of each jump. To obtain population averaged poses for the jump start-, mid- and end-points, we normalized each pose equivalently to the analysis of gait data. This aligned the resulting poses at their origin and we were able to calculate characteristic jump poses by averaging them across the entire population. For the population averaged mean angle traces we aligned each individual trace according to the mid-point of each jump and then averaged across the entire population. To highlight the diversity of the data given by the reconstructed poses, we calculated distance- and auto-correlations of several different metrics and joints: jump distances were correlated with spatial z-velocities and angular velocities of spine joints at time points up to 400 ms before the end of a jump and absolute spatial velocities and angular velocities of hind limbs joints were correlated with each other at the start-point of a jump. Since differences in bone lengths for each animal dominated the correlation for spatial position and joint angle we only focused on their first temporal derivatives.

### Computing hardware

All pose reconstructions and analyzes were conducted on a workstation equipped with an AMD Ryzen 7 2700x CPU, 32 GB DDR4 RAM, Samsung 970 EVO 500 GB SSD, and a single NVIDIA GeForce RTX 1080 Ti (11 GB) GPU. The installed operating system was Ubuntu 18.04.5 LTS. Training DeepLabCut was either conducted on a NVIDIA GeForce RTX 1080 Ti (11 GB) GPU, using CUDA version 10.0 and NVIDIA driver version 410.48, or a NVIDIA GeForce RTX 2080 Ti (11 GB) GPU, using CUDA version 11.0 and NVIDIA driver version 450.80.02.

## Code availability

Code for performing pose reconstructions will be made publicly available on GitHub: https://github.com/bbo-lab/ACM

## Data availability

Raw data available on request.

---

**Supplementary Fig. 1.**
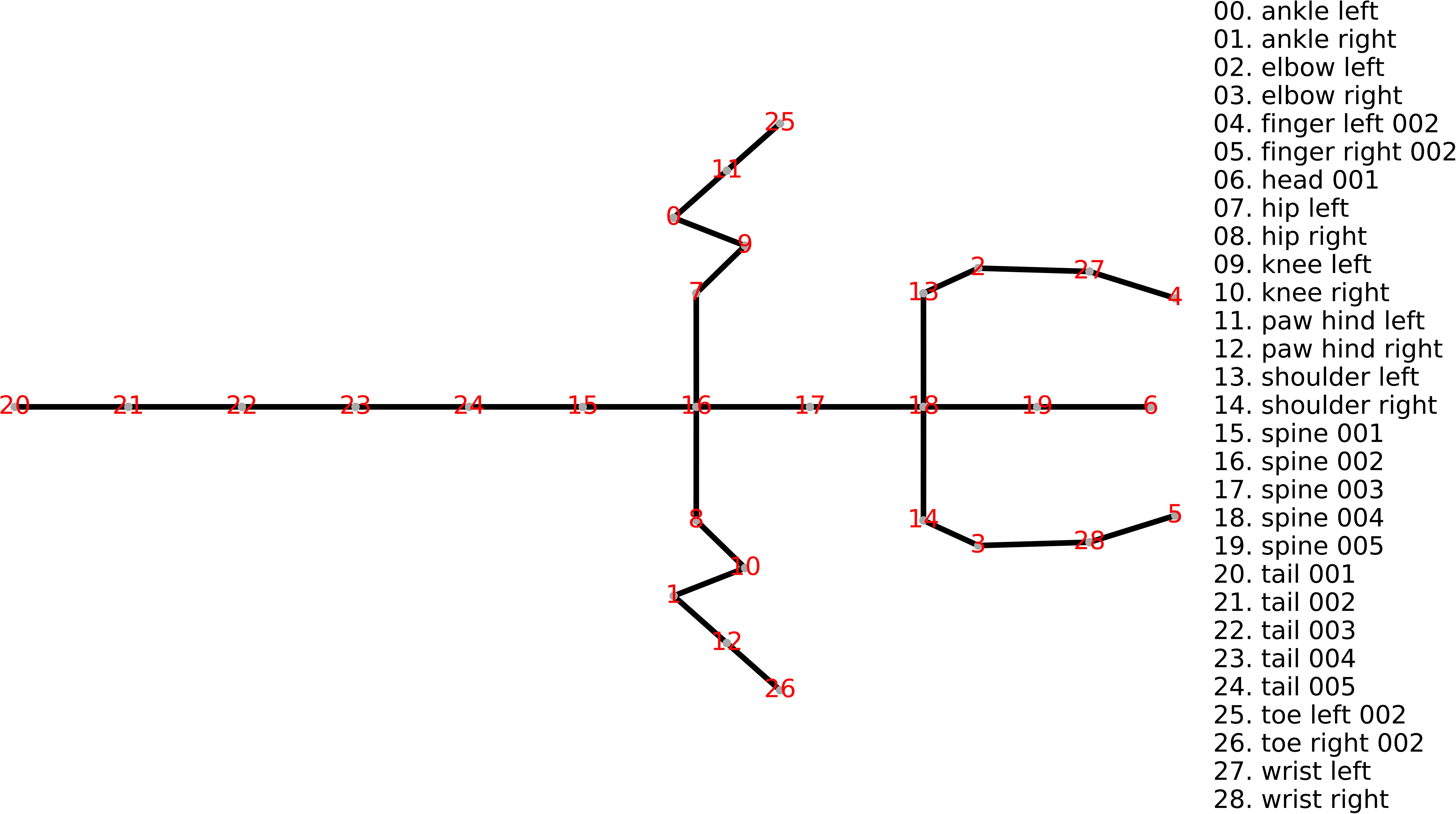
Projection of the generalized skeleton model in the xy-plane with index and name for each individual joint. For this figure, all bone lengths are equal and all bone rotations have been set to the mean of their upper and lower bounds.

**Supplementary Fig. 2.**
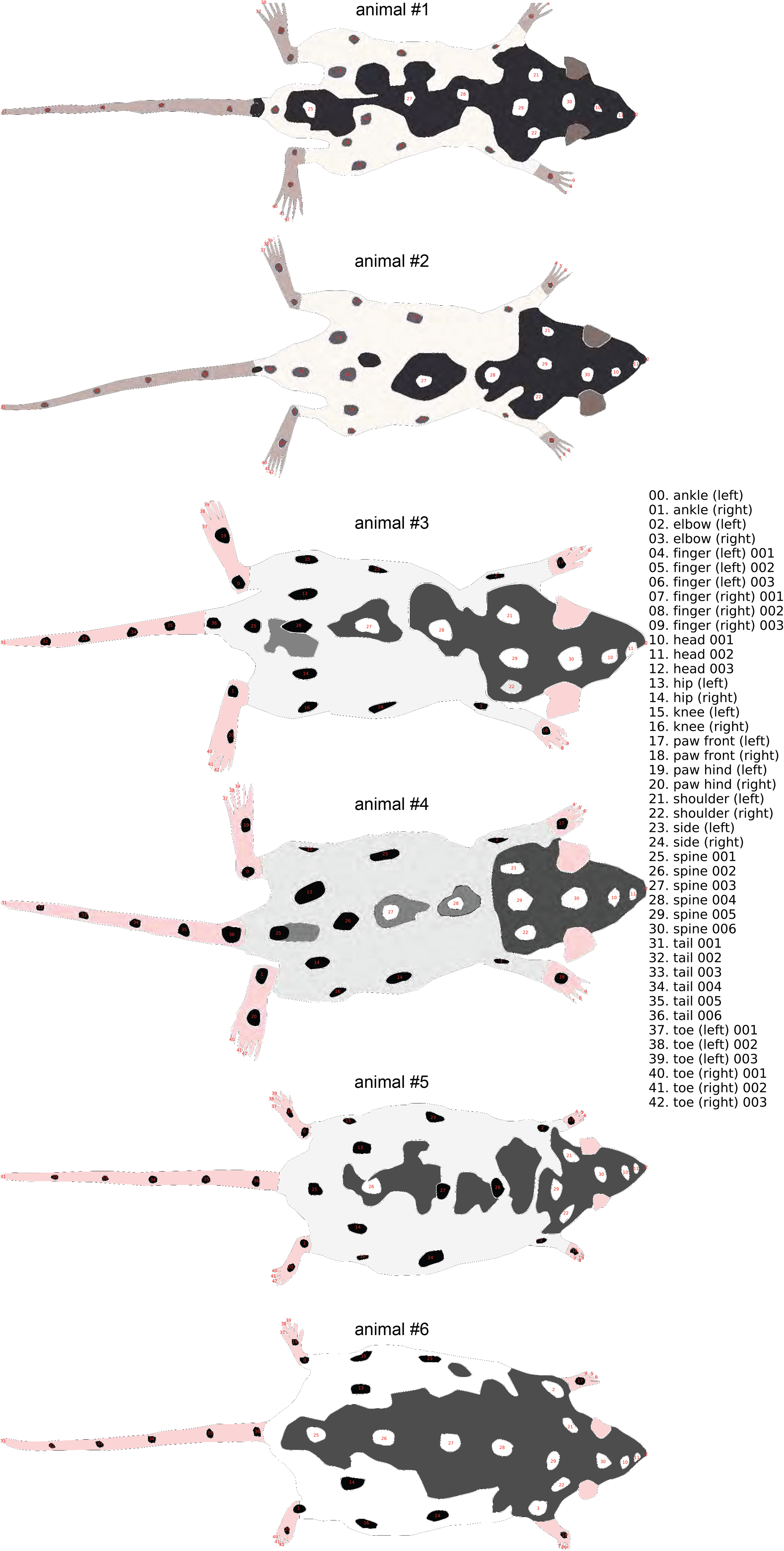
Schematic images of the six labeled animals with index and name for each individual surface marker

**Supplementary Fig. 3.**
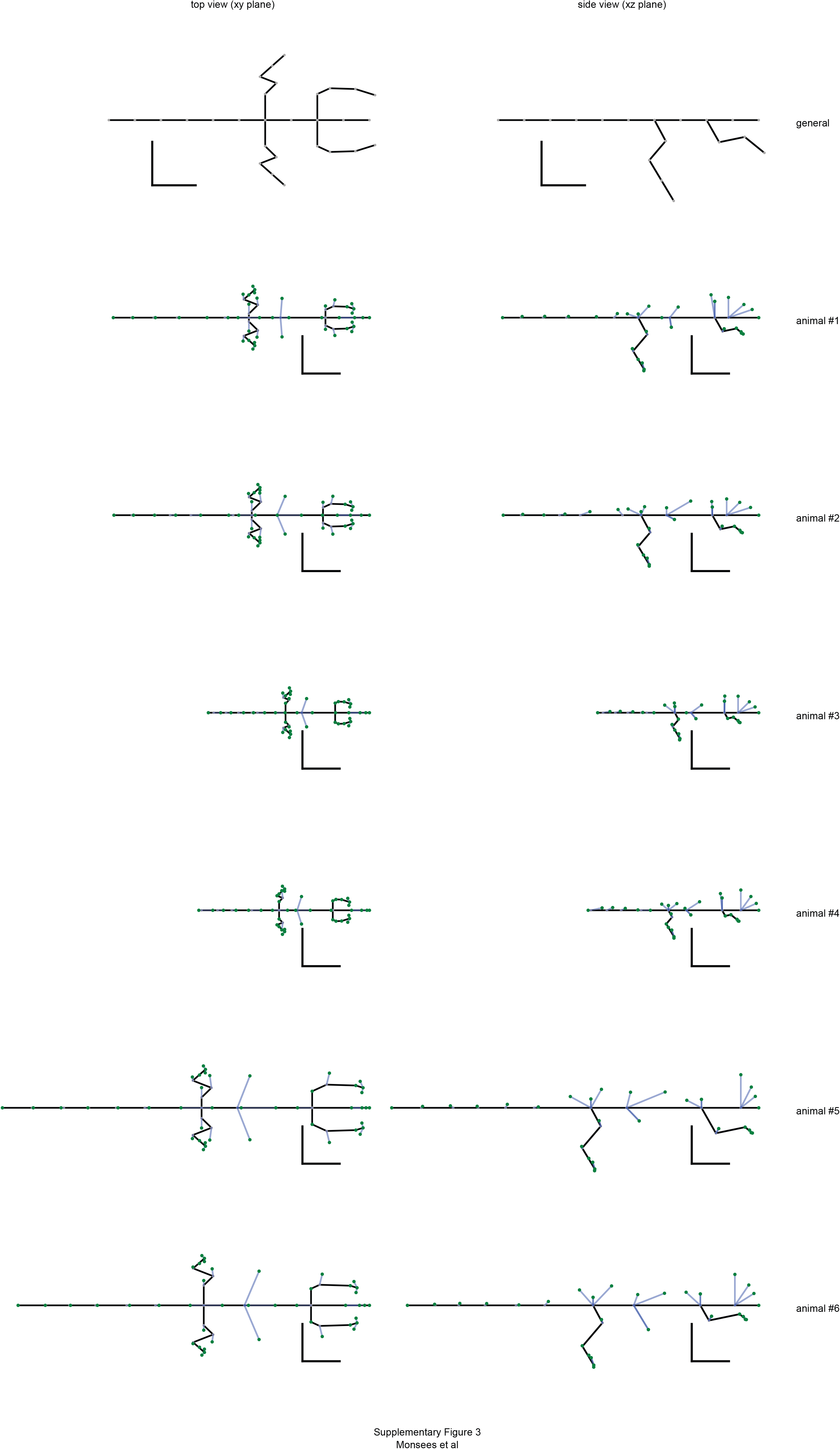
Projections of the generalized and learned skeleton models for each animal as viewed from the top (xy-view, left column) and from the side (xz-view, right column). All bone rotations were set to the mean of their upper and lower bounds. Green dots indicate the learned positions of surface markers and blue lines join paired joints and surface markers. All scale bars 5 cm.

**Supplementary Fig. 4.**
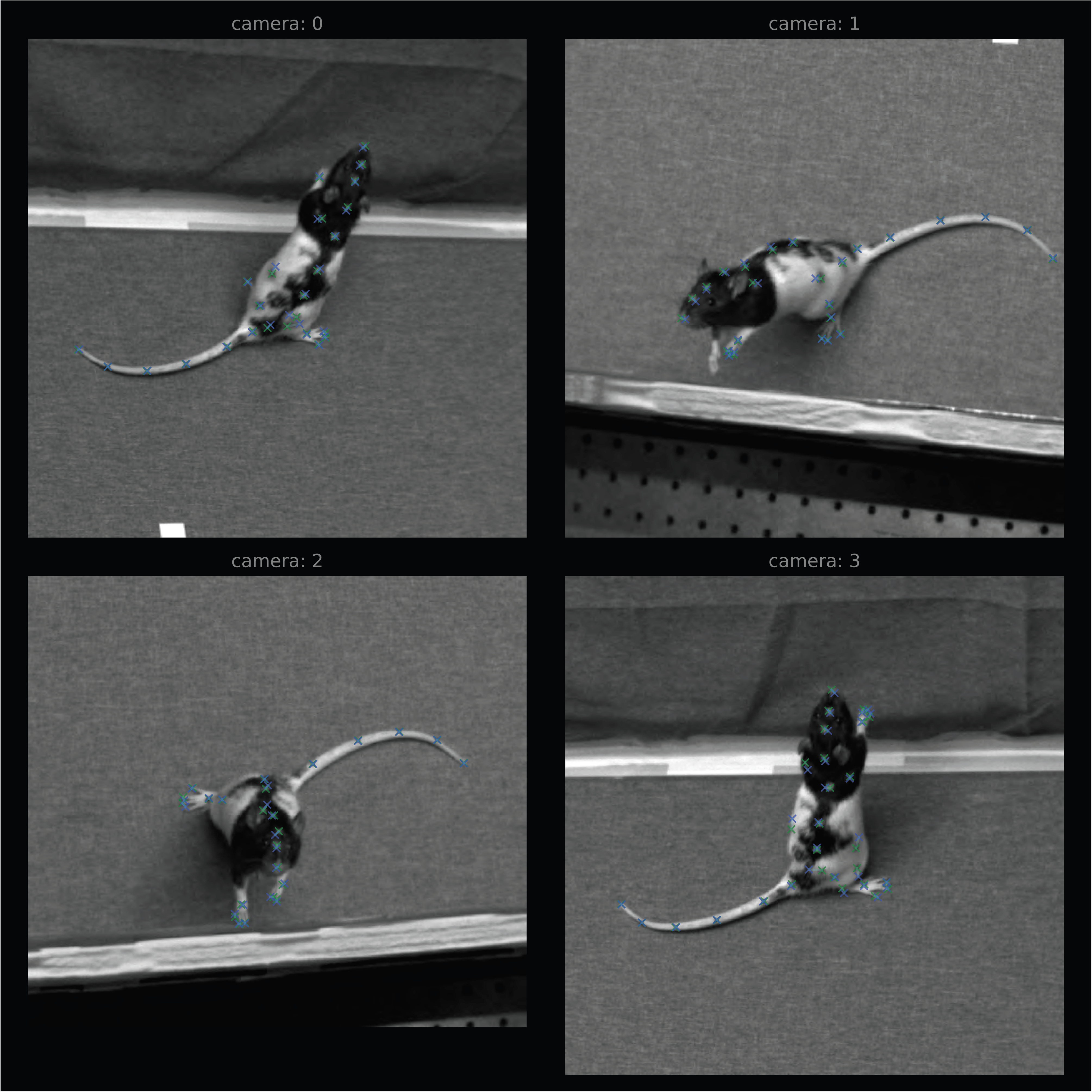
Synchronous training frames from four calibrated cameras used for learning skeleton lengths and surface maker positions. The figure shows the manually labeled surface marker positions (green), their locations after the skeleton model is learned (blue) and the discrepancies between the two (green lines).

**Supplementary Fig. 5.**
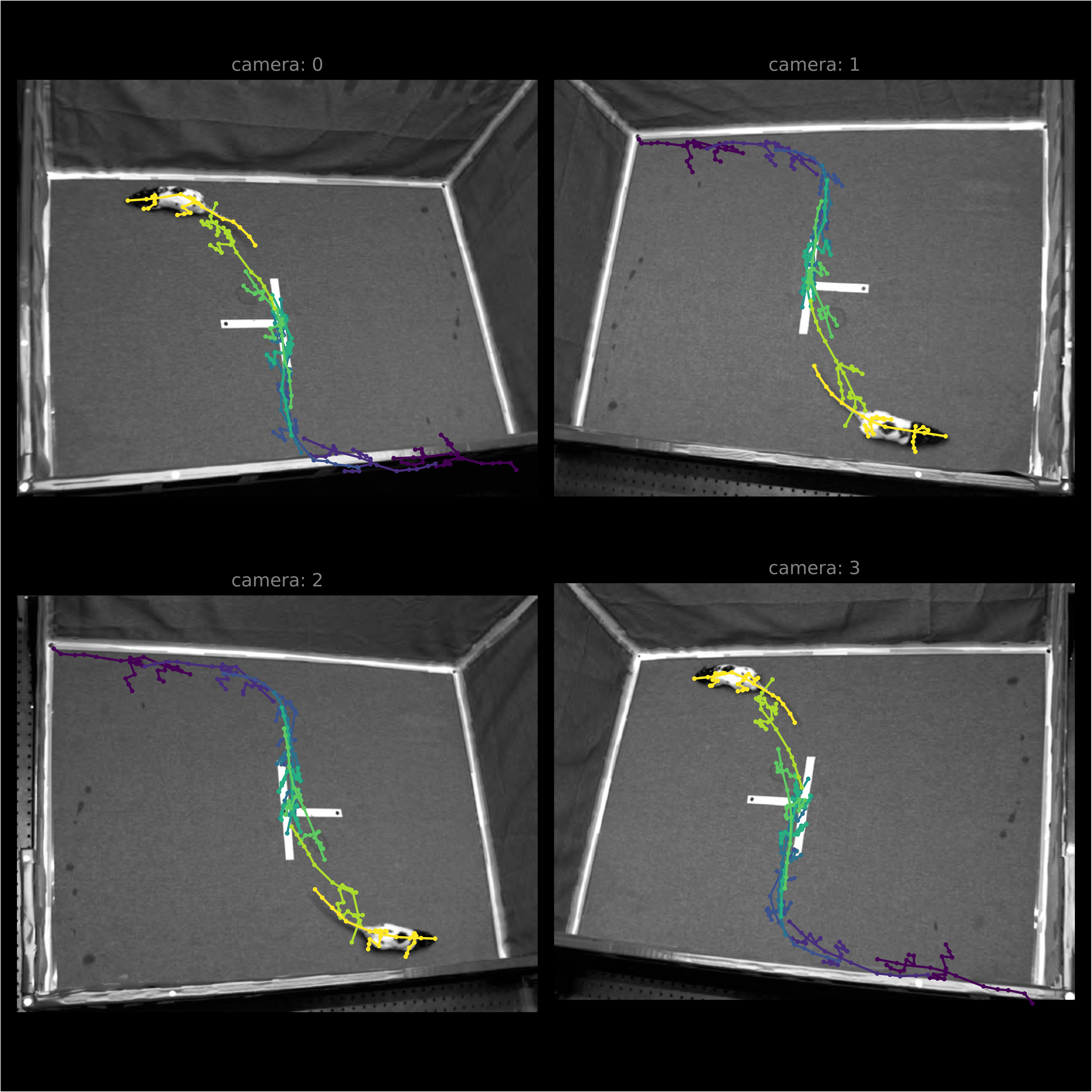
Synchronous frames from four calibrated cameras which were part of the gait data set. Reconstructed skeleton poses are shown for different time points of the gait sequence. The time difference between poses is 1 s.

**Supplementary Fig. 6.**
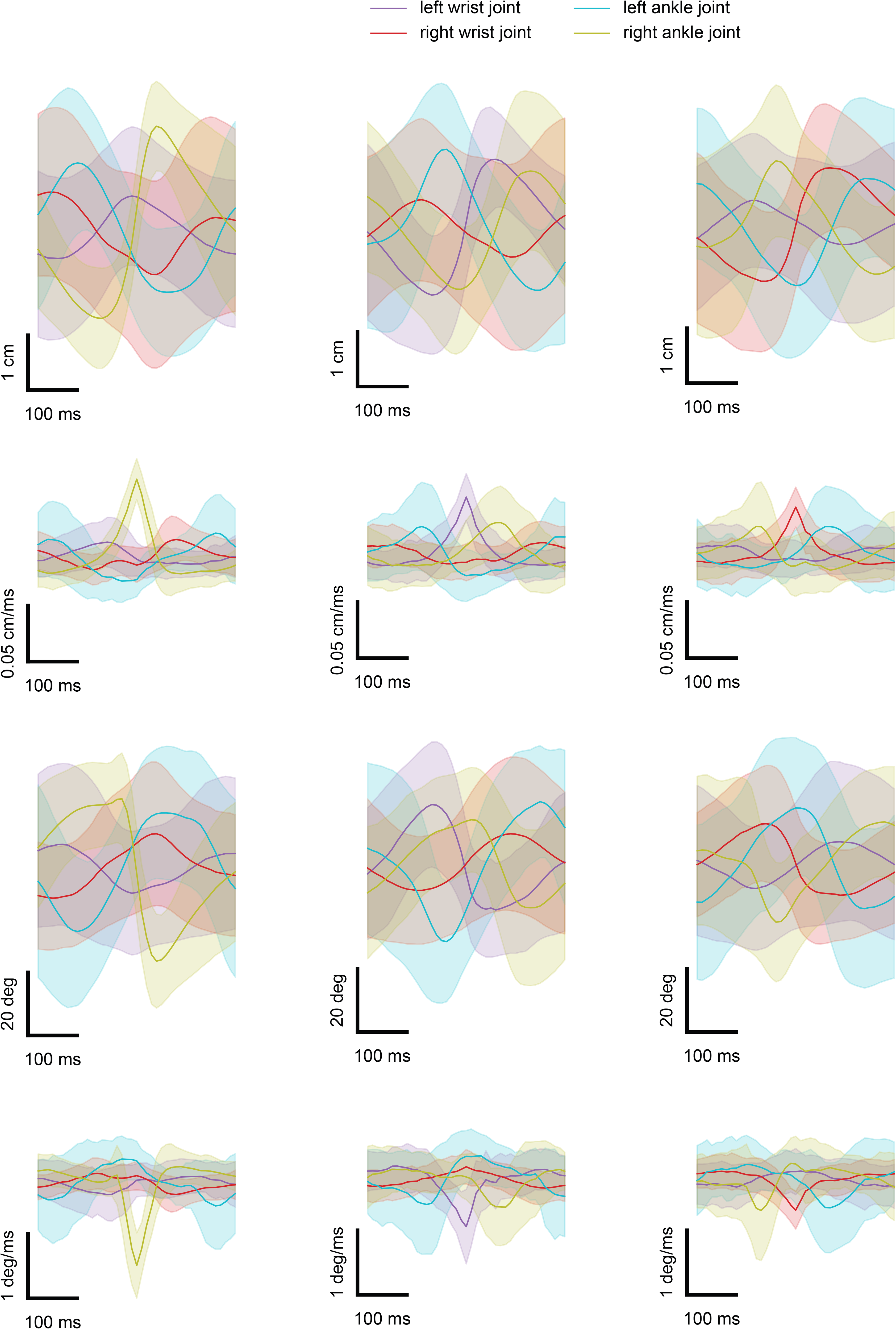
Averaged traces from the ACM as in Figure 3d,g,j,m (left), but with trace alignment based on the velocity peaks for the right ankle (right column), left wrist (center column) and right wrist joint (right column), instead of aligning to the velocity peak of the left ankle joint.

**Supplementary Fig. 7.**
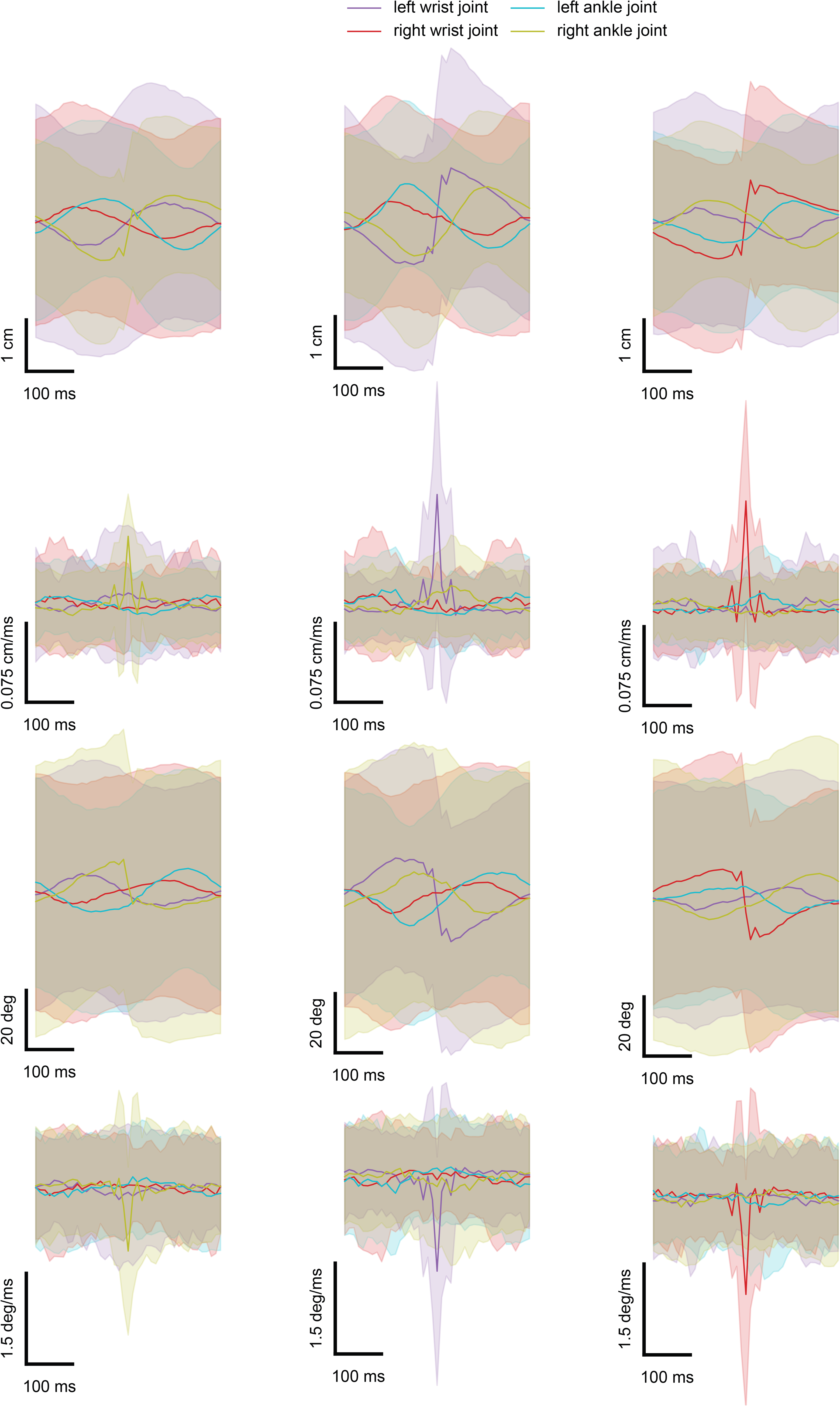
Averaged traces from the model not constrained by either temporal or joint limit constraints as in Figure 3d,g,j,m (right) aligned to velocity peaks for the right ankle (right column), left wrist (center column) and right wrist joint (right column), instead of aligning to the velocity peak of the left ankle joint.

**Supplementary Fig. 8.**
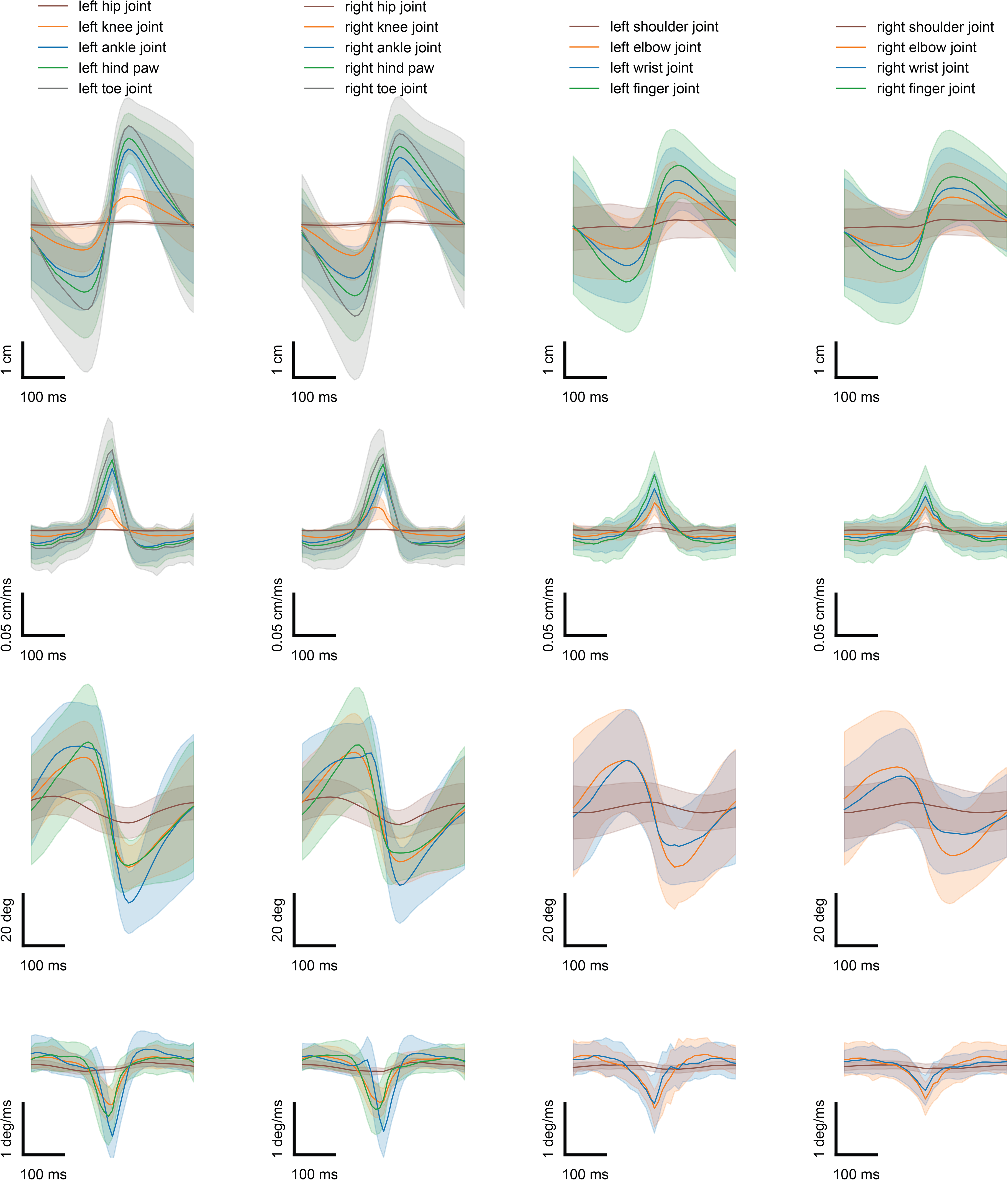
Averaged traces from the ACM for all joints of the left hind (left column), right hind (center left column), left front (center right column) and right front limb (right column). Traces were aligned to velocity peaks from the left ankle, right ankle, left wrist and right wrist joint of the respective limb.

**Supplementary Fig. 9.**
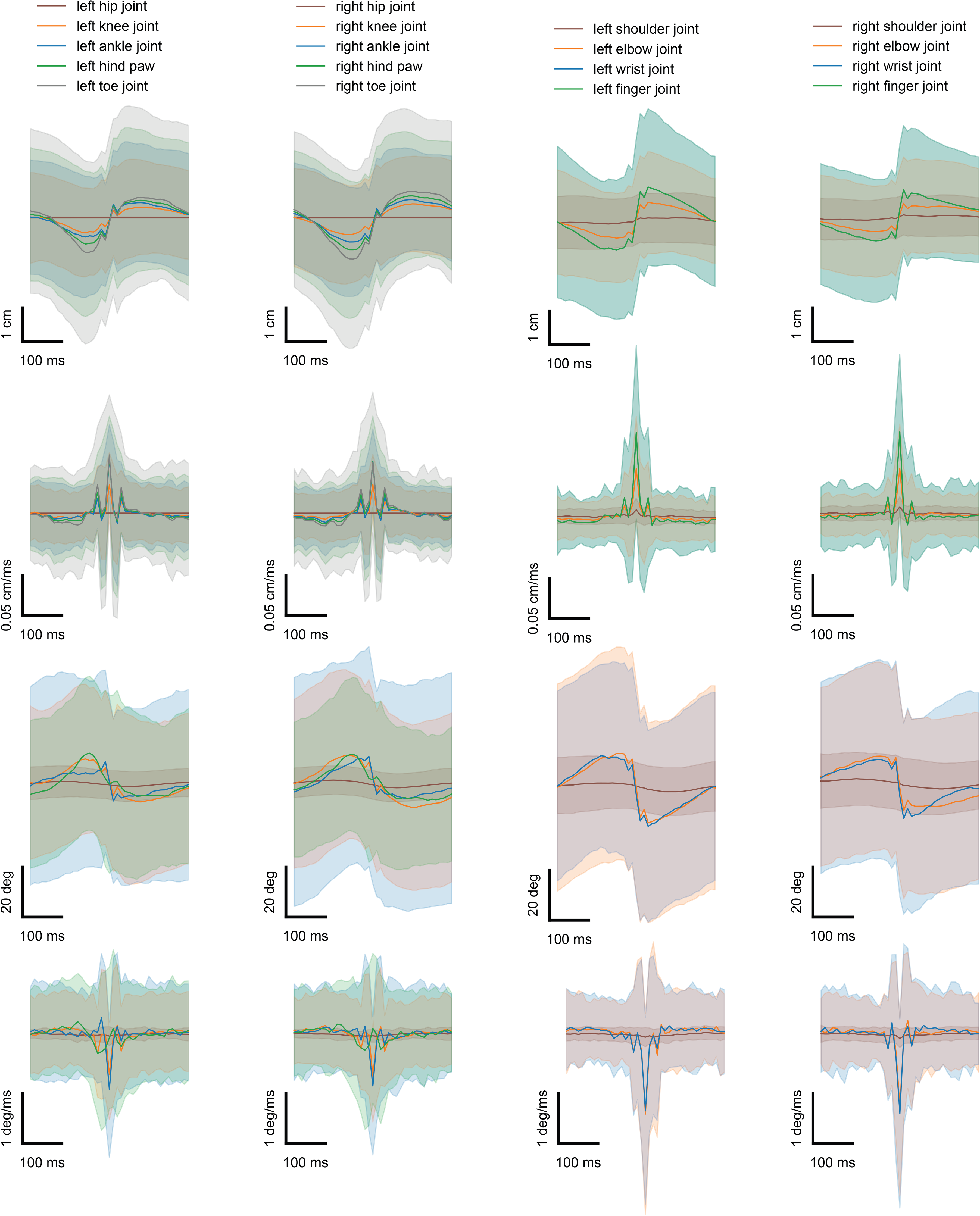
Averaged traces from the model not constrained by either temporal or joint limit constraints for all joints of just the left hind (left column), right hind (center left column), left front (center right column) and right front limb (right column). Traces were aligned to velocity peaks from the left ankle, right ankle, left wrist and right wrist joint of the respective limb.

**Supplementary Fig. 10.**
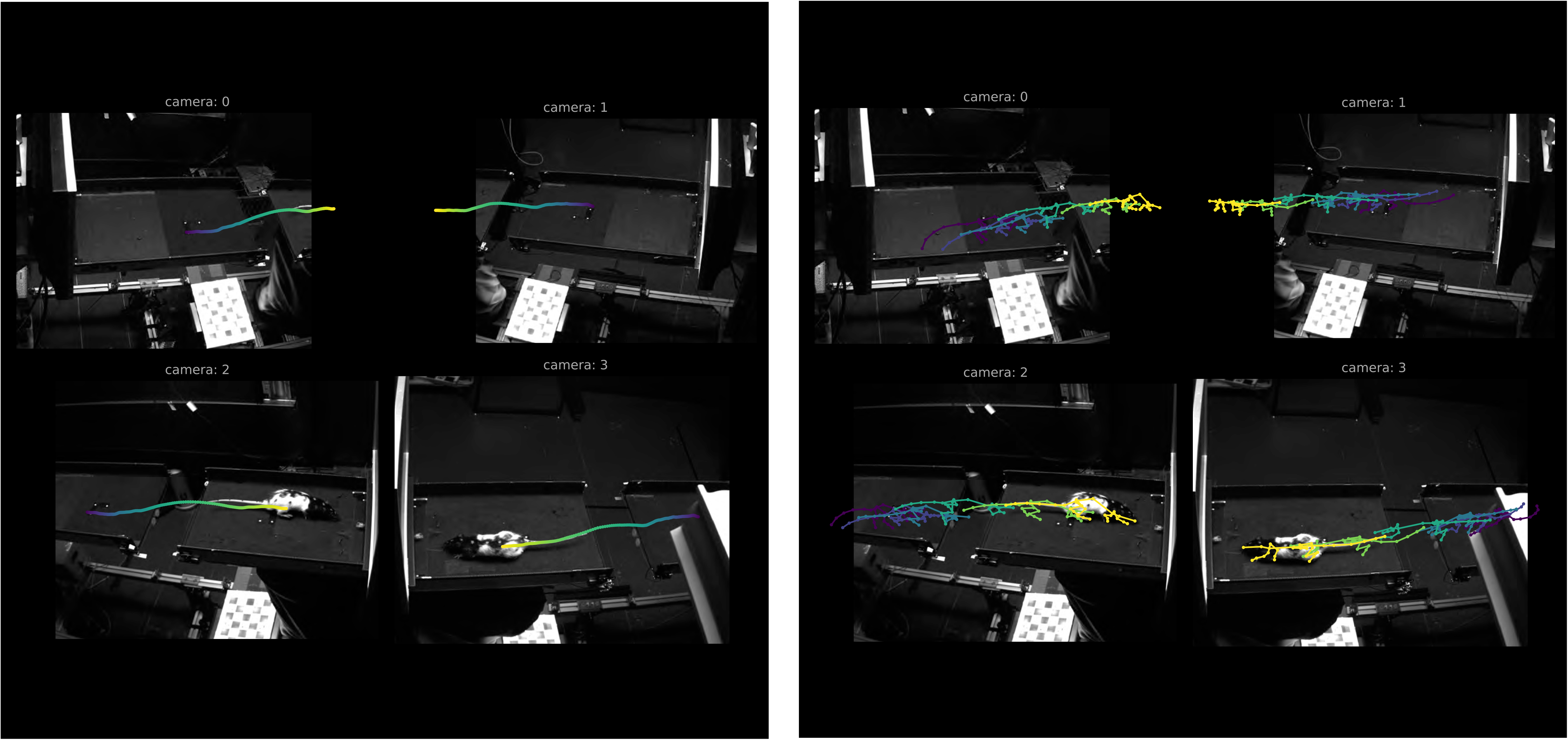
Synchronous frames from four calibrated cameras which were part of the gap-crossing data set, with overlay of the center of mass (left) and reconstructed skeleton poses (right) shown for different time points.

**Supplementary Fig. 11.**
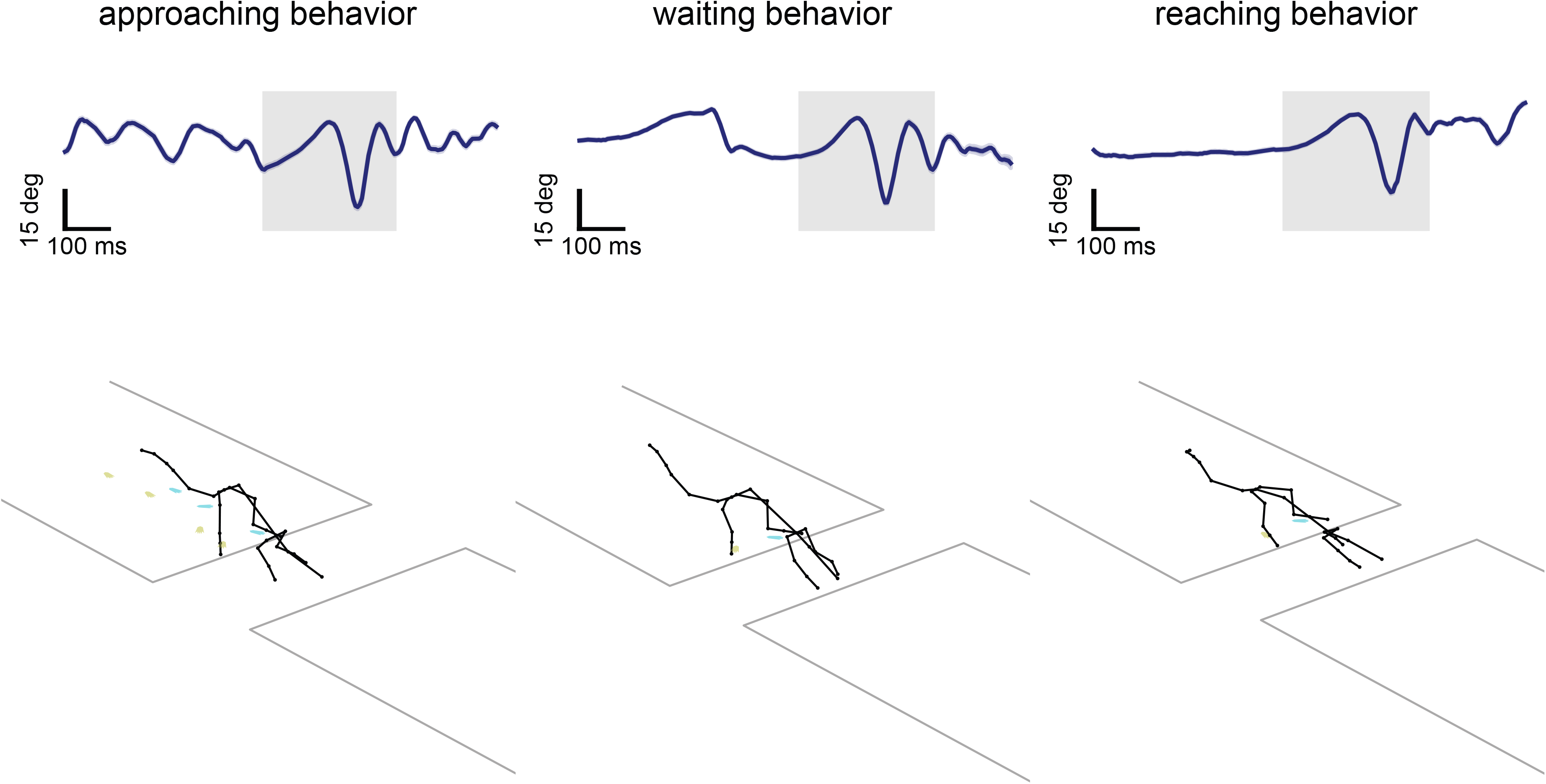
Averaged joint-angle trace (average angles of all spine and hind paw joints) as a function of time (top) and reconstructed poses (bottom), showing hind paw footprints (cyan/yellow), at the start of a jump for three different gap crossing events. The animal was approaching the gap fast, with a smooth transition from walking to jumping behavior (left), waiting at the edge (center) or reaching to the other side of the track with its right front limb (right) before crossing the gap.

**Supplementary Fig. 12.**
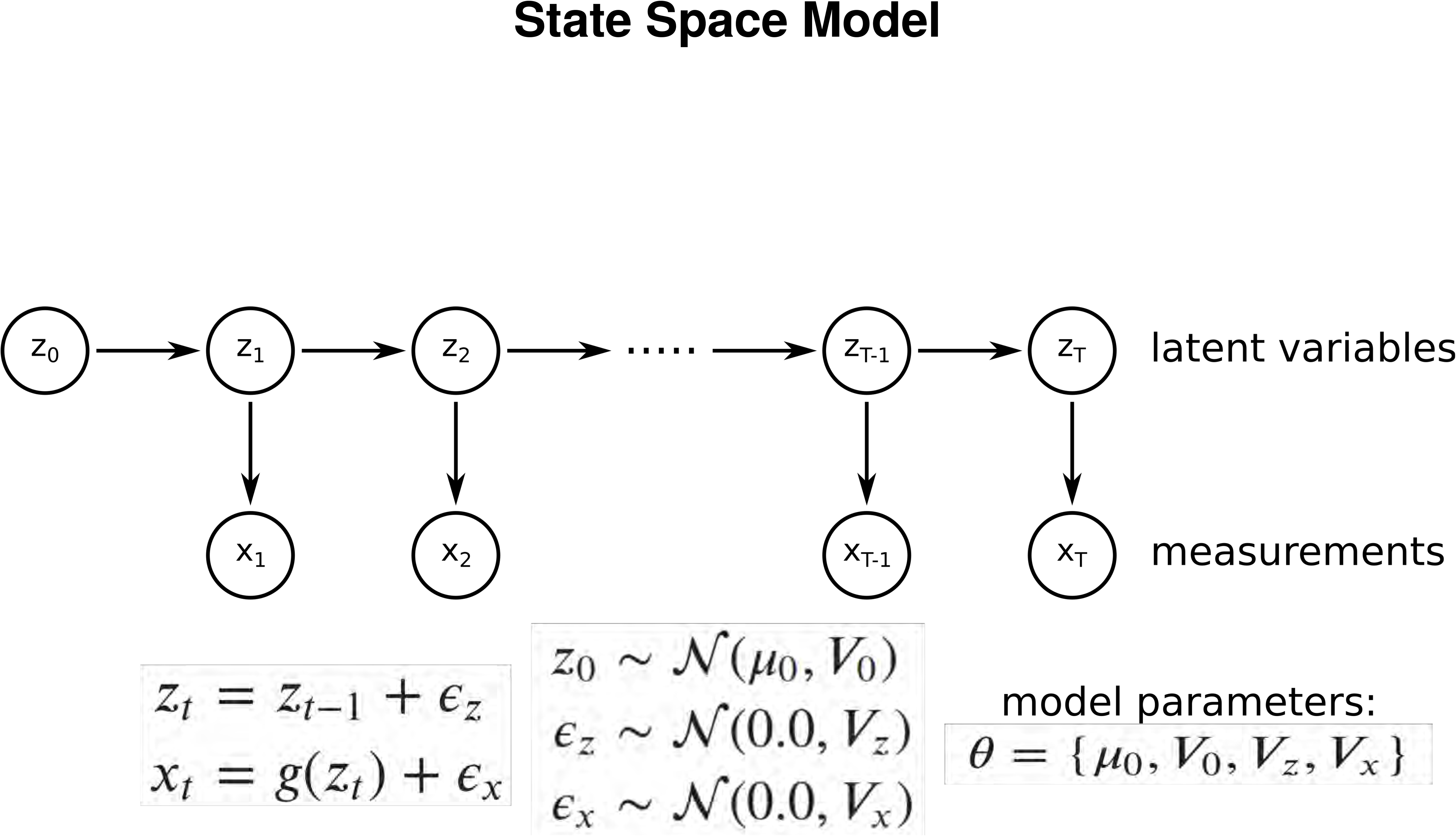
Illustration of the state space model used for describing behavioral time series’.

## 1 Parameterizing rotations

We choose to parameterize rotations with Rodrigues vectors as they are well suited for the description of bone rotations with three rotational degrees of freedom [10]. A Rodrigues vector *r* is formed by combining the axis of rotation *ω* ∈ ℝ^3^ and the rotation angle *θ* ∈ ℝ:

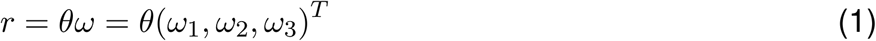

where ||*ω*|| = 1. To calculate the associated rotation matrix *R* from a given Rodrigues vector *r* we can use the following function:

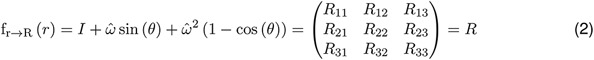

where *I* ∈ ℝ^3×3^ is the identity matrix and 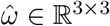 is given by:

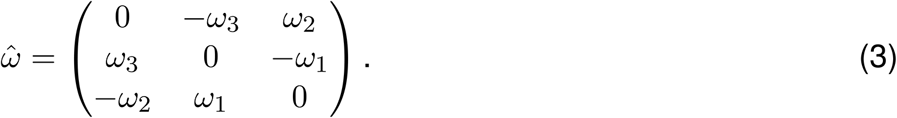

## 2 Camera calibration

### 2.1 Pinhole camera model

To project an arbitrary three-dimensional joint or surface marker location *m*_3D_ ∈ ℝ^3^ onto a camera sensor to obtain the corresponding two-dimensional data point *m*_2D_ ∈ ℝ^2^, we are using a pinhole camera model [6], which gives the following relationship between the two:

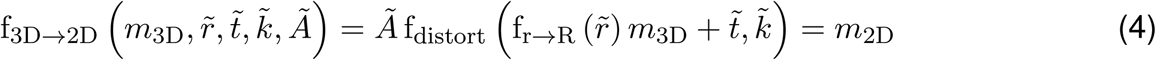

where 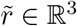 is the Rodrigues vector and 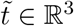 the translation vector of the respective camera, such that the expression 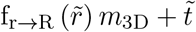 maps *m*_3D_ from the world coordinate system into the coordinate system of the camera. Given the camera’s distortion vector 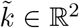, the function f_distort_ applies radial distortions according to

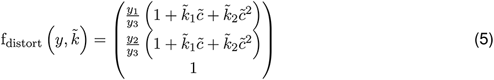

with *y* = (*y*_1_, *y*_2_, *y*_3_)^*T*^ and 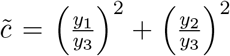. The final mapping onto the two-dimensional camera sensor is done using the camera matrix 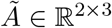 given by

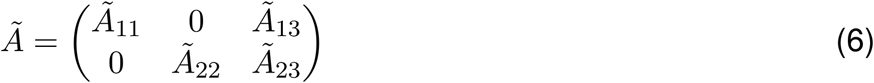

where 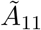 and 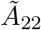 are the focal lengths and 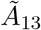 and 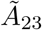 are the x- and y-location of the camera’s optical center.

### 2.2 Calibration of multiple cameras

Given a multi-camera setup with several cameras and overlapping fields of view, we need to infer the initially unknown location and camera parameters of every individual camera in the setup as this allows us to predict where a three-dimensional point in space will be visible on each camera sensor. This can be achieved by generating a sequence of images showing an object whose physical structure and dimensions are known to us. Hereby, the images are taken synchronously in all cameras, such that the spatial location and orientation of the shown object is identical for a given set of images at a certain time point. For this purpose checkerboards are suited objects as edges of individual tiles can be detected automatically in recorded image frames and the description of their spatial structure requires only a single parameter, i.e. the length of a quadratic tile. Given a multi-camera setup with *n*_cam_ cameras and *n*_time_ time points at which we used each camera to record images, which show a checkerboard that has a total of *n*_edge_ detectable edges, we can calibrate the setup by minimizing a respective objective function via gradient decent optimization using the Trust Region Reflective algorithm [2]:

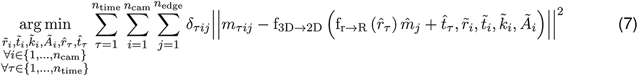

where 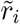 is the Rodrigues vector, 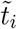 is the translation vector, 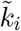 is the distortion vector and 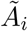 is the camera matrix of camera *i*. The Rodrigues vector 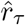 and the translation vector 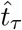 encode the orientation and translation of the checkerboard at time point *τ*. Since the checkerboard is a planar object each edge *j* is given by a three-dimensional point 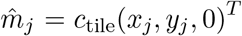 with the known length of a single tile *c*_tile_ and *x_j_* ∈ ℕ as well as *y_j_* ∈ ℕ. Furthermore, the two-dimensional edge *j* in camera at time point *τ* is denoted as *m_τij_* ∈ ℝ^2^ and the delta function *δ_τij_* indicates whether this edge is detected successfully, i.e. *δ_τij_* = 1, or not, i.e. *δ_τij_* = 0.

## 3 Skeleton model

### 3.1 Modifying the skeleton model to obtain new poses

Given a three-dimensional skeleton model, we need to adjust joint locations by rotating each bone of the model, such that resulting three-dimensional positions of rigidly attached surface markers match the respective two-dimensional locations in our video data. Assuming our skeleton model has a total of *n*_bone_ bones and *n*_marker_ surface markers, we want to generate the three-dimensional locations of the joints 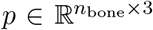 and surface markers 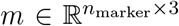, which can be obtained according to Algorithm 1.

**Algorithm 1.**
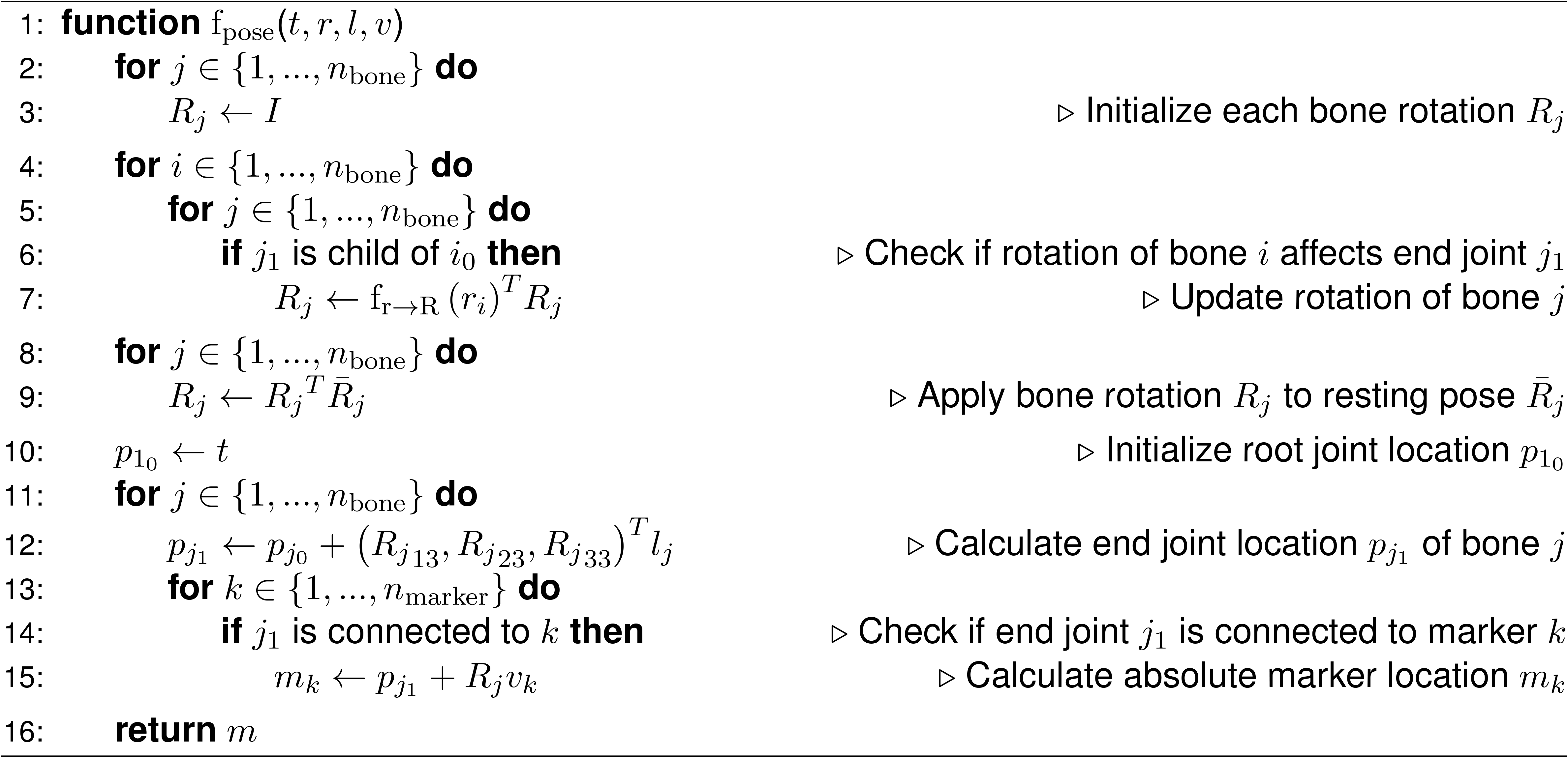

Here, it is assumed that the set {1, …, *n*_bone_} is sorted, such that one iterates through the skeleton graph beginning with the bone whose start joint is the root joint 1_0_ and then proceed with the bones further down the skeleton graph. Thus, it is always guaranteed that for *j* > *i*, the start joint *i*_0_ of bone *i* is never a child of the start joint *j*_0_ of bone *j*. It is also assumed that the bone coordinate systems of the skeleton model are constructed such that their z-directions encode the directions in which the respective bones are pointing. Furthermore, the global translation vector *t* ∈ ℝ^3^ corresponds to the three-dimensional location of the skeleton’s root joint, the rows of the tensor 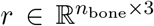 contain Rodrigues vectors encoding the bone rotations, the vector 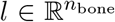 contains the bone lengths and the rows of the tensor 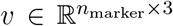 contain the relative maker locations, i.e. the locations of the markers when the position of the attached joints are assumed to be the origin. The resting pose 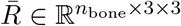 of the animal describes the orientation of the bones when no additional rotations are applied, i.e. *r_i_* = (0, 0, 0)^*T*^ ∀ *i* ∈ {1, …, *n*_bone_}. Here, the frequent usage of the transpose operation allows to first rotate bones, which are the closest to the leaf joints of the skeleton graph [4]. This has the advantage that we can enforce constraints on bone rotations with reference to a global coordinate system that corresponds to the three main axes of the animal’s body. Assume we only model a single front limb where we only have rotations around the shoulder, elbow and wrist, i.e. *R*_shoulder_, *R*_elbow_ and *R*_wrist_, and would like to obtain the new orientation *R*_new_ of the bone whose start joint is identical to the animal’s wrist given its resting pose 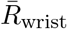 while iterating through the skeleton graph starting from the root joint, i.e. the shoulder. Then we can obtain *R*_new_ according to

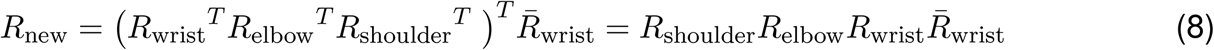

Thus, we can iterate through the skeleton graph from the root to the leaf joints but actually apply the respective bone rotations in the reversed order.

### 3.2 Inferring bone lengths and surface marker positions

Reconstructing poses for *n*_time_ time points can be archived equivalently to the calibration of a multi-camera setup as discussed in Section 2.2, i.e. we need to minimize a respective objective function via gradient decent optimization using the L-BFGS-B algorithm [3]:

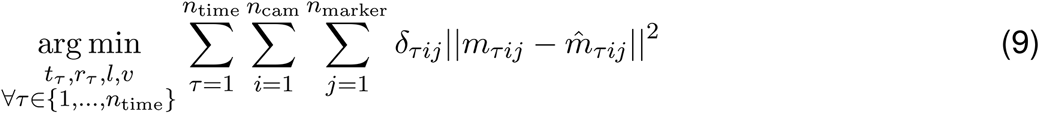

where *m_τij_* is the two-dimensional location of marker *j* in camera *i* at time point *τ* and *δ_τij_* indicates whether this marker location was successfully detected, i.e. *δ_τij_* = 1, or not, i.e. *δ_τij_* = 0. The corresponding projected two-dimensional marker location 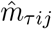 can be obtained by propagating the absolute marker positions calculated via Algorithm 1 through the projection function f_3D→2D_:

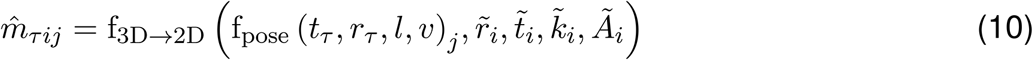

where *t_τ_* ∈ ℝ^3^ and 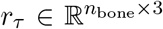 denote the translation vector and the bone rotations at time point *τ*. Note how there is a set of pose-encoding parameters *t_τ_* and *r_τ_* for each time point *τ* whereas the bone lengths *l* and the relative surface marker positions *v*, which encode the animal’s skeletal structure and configuration, are shared across all time points. Thus, if we provide enough time points where the animal is visible in many different poses, which ideally cover the entire spectrum of the animal’s behavioral space, we can not only reconstruct the pose of the animal for the given time points but are also able to learn the structure of the animal’s skeleton, by inferring the unknown parameters *l* and *v*.

### 3.3 Scaling of input and output variables

In general, we always scale the translation vector *t* and the bone rotations *r* as well as the resulting two-dimensional marker locations 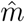, such that all of them roughly lie within the same range, i.e. [−1, 1]. Particularly, we define the normalization constants *c_t_* = 50 cm and 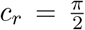 rad as well as *c*_1_ = 640 px and *c*_2_ = 512 px, which we use to normalize *r* and *t* as well as 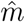. The choice for *c_t_* was based on the dimensions of the largest arena we used in our experiments, where the maximum distance to an arena’s edge from the origin of the world coordinate system, located at the center of the arena, was around 50 cm. The choice for *c_r_* was based on the maximum bone rotation of the naively constrained spine and tail joints in our skeleton model, which was equal to 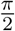 rad. The choice for *c*_1_ and *c*_2_ were based on the sensor sizes of the cameras we used in our experiments, which were all equal to 1280 × 1024 px^2^. Using the normalization constants we obtain the normalized translation vector 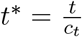 and the normalized bone rotations 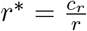 as well as the normalized two-dimensional marker locations

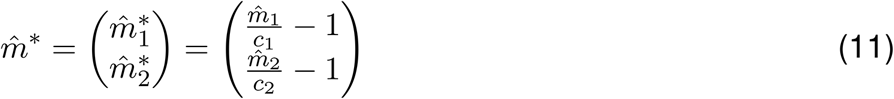

for a single two dimensional marker location 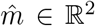, such that 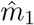 represents its x- and 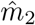 its y-coordinate. These normalized variables were used instead of their non-normalized counterparts in all depicted optimization and pose reconstruction steps.

### 3.4 Enforcing body symmetry

To improve the inference of bone lengths and surface marker positions we took advantage of the symmetric properties of an animal’s body, i.e. for every left-sided limb there exists a corresponding limb on the right side. Furthermore, we also placed the surface markers onto the animal’s fur, such that the marker-pattern itself was symmetrical, e.g. for a marker that was placed to a position close to the left hip joint there was a corresponding marker on the right side of the animal. By incorporating this knowledge into Algorithm 1 we reduced the number of free parameters, i.e. we only optimized the reduced bone lengths 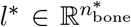 and relative marker positions 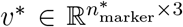, where 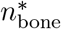 is the number of asymmetrical bones, i.e. bones along the head, spine and tail, plus the number of limb bones on the animal’s left side and, equivalently, 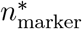 denotes the number of the asymmetrical and left-sided markers. The excluded right-sided limb bones were then enforced to have the same lengths as the corresponding limb bones on the left side. Additionally, we also applied this concept for the relative marker locations by mirroring the x-component of the left-sided markers at the yz-plane to obtain the relative marker locations of the markers on the right side. To implement this we defined Algorithm 2, which maps the reduced bone lengths *l*^∗^ to the original parameter *l*.

**Algorithm 2.**
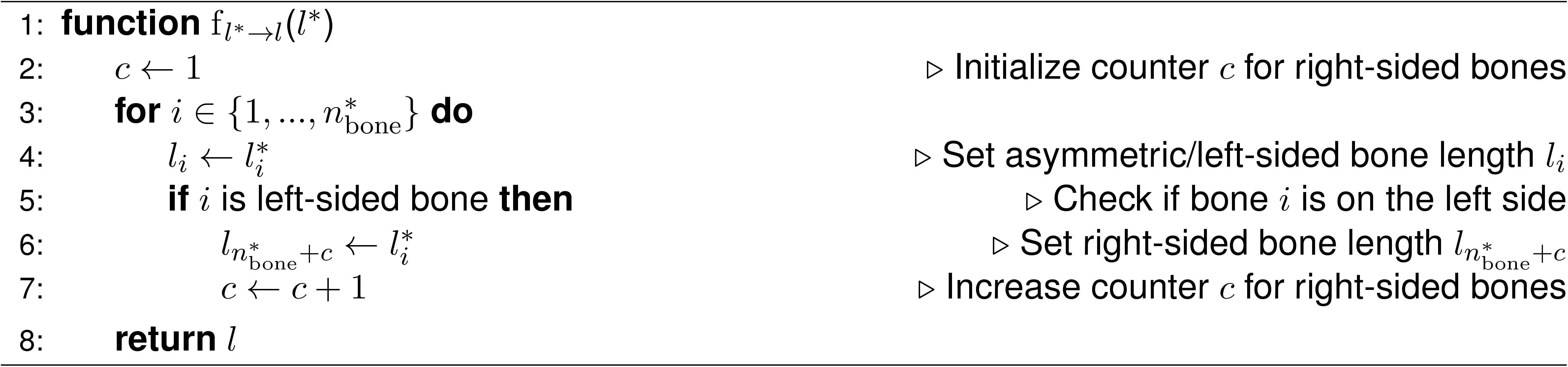

Equivalently, we also defined the corresponding Algorithm 3, which maps the reduced relative marker positions *v*^∗^ to their original counterpart *v*.

**Algorithm 3.**
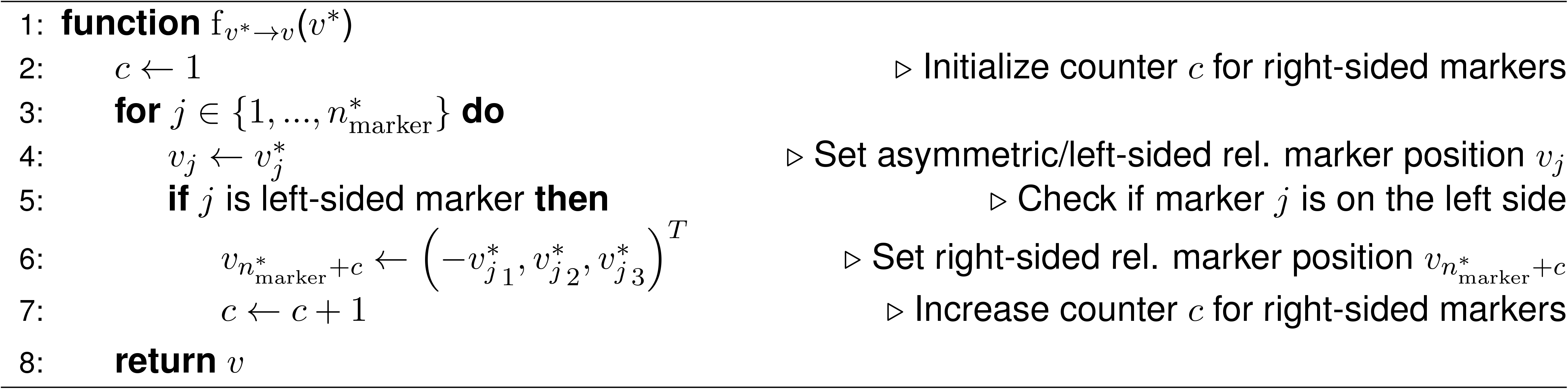

To learn the underlying three-dimensional skeleton model while also enforcing body symmetry, we then redefined 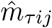 from equation 10 as follows:

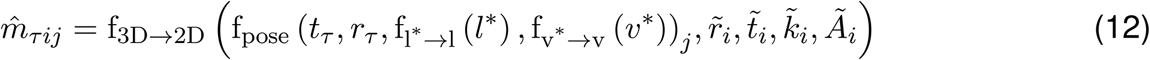

and minimized equation 9 with respect to the parameters *l*^∗^ and *v*^∗^ instead of *l* and *v*.

## 4 Probabilistic pose estimation

### 4.1 Using a state space model to describe behavioral time series’

To allow for probabilistic pose reconstruction of entire behavioral sequences of length *T*, which ensures that poses of consecutive time points are similar to each other, we deploy a state space model, given by a transition and an emission equation

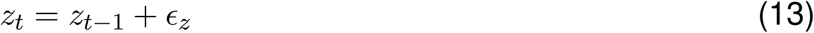

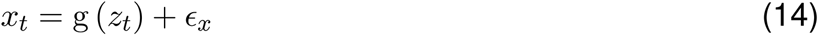

where at time point *t* ∈ {1, …, *T*} the state variable 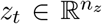 encodes the position of the animal as well as the bone rotations and the measurement variable 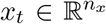 represents the two-dimensional surface marker locations in all cameras given by a trained neural network. Thus, the state variable *z*_*t*_ contains the global translation vector *t* as well as the pose-encoding tensor *r* for time point *t* and the measurement variable *x_t_* is a constant quantity given for all time points *t*. The function g, given by Algorithm 4, computes the noise-free measurements of the two-dimensional surface marker locaions 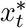 given the state variable *z_t_*. At this point the bone lengths *l* and relative maker locations *v* are already inferred and therefore given. The same applies to the Rodrigues vector 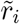, the translation vector 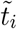, the distortion vector 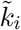 and the camera matrix 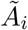 of camera *i*, which we obtained from calibrating the multi-camera setup. The normalization constants *c_t_*, *c_r_* as well as *c*_1_ and *c*_2_ are the same as in Section 3.3. The probabilistic nature of the model is given by incorporating the two normally distributed random variables 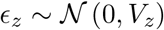 and 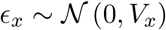, simulating small pose changes over time and measurement noise, as well as the initial state 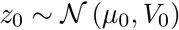, which is also assumed to be a normally distributed random variable. Thus, the state space model is entirely described by the model parameters 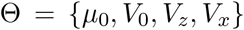. This allows for inferring a set of expected state variables 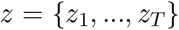 given our measurements 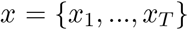 in case we have a good estimate for the model parameters Θ. Alternatively, we are also able to calculate a set of model parameters Θ, which, given an estimate for the state variables *z*, maximizes a lower bound of the model’s evidence, i.e. the evidence lower bound (ELBO). The former is equivalent to the expectation step (E-step) of the expectation-maximization (EM) algorithm, which can be performed by applying the unscented Rauch-Tung-Striebel (RTS) smoother, whereas the latter is identical to the algorithm’s maximizaion step (M-step), in which new model parameters are calculated in closed form to maximize the ELBO [8].

**Algorithm 4.**
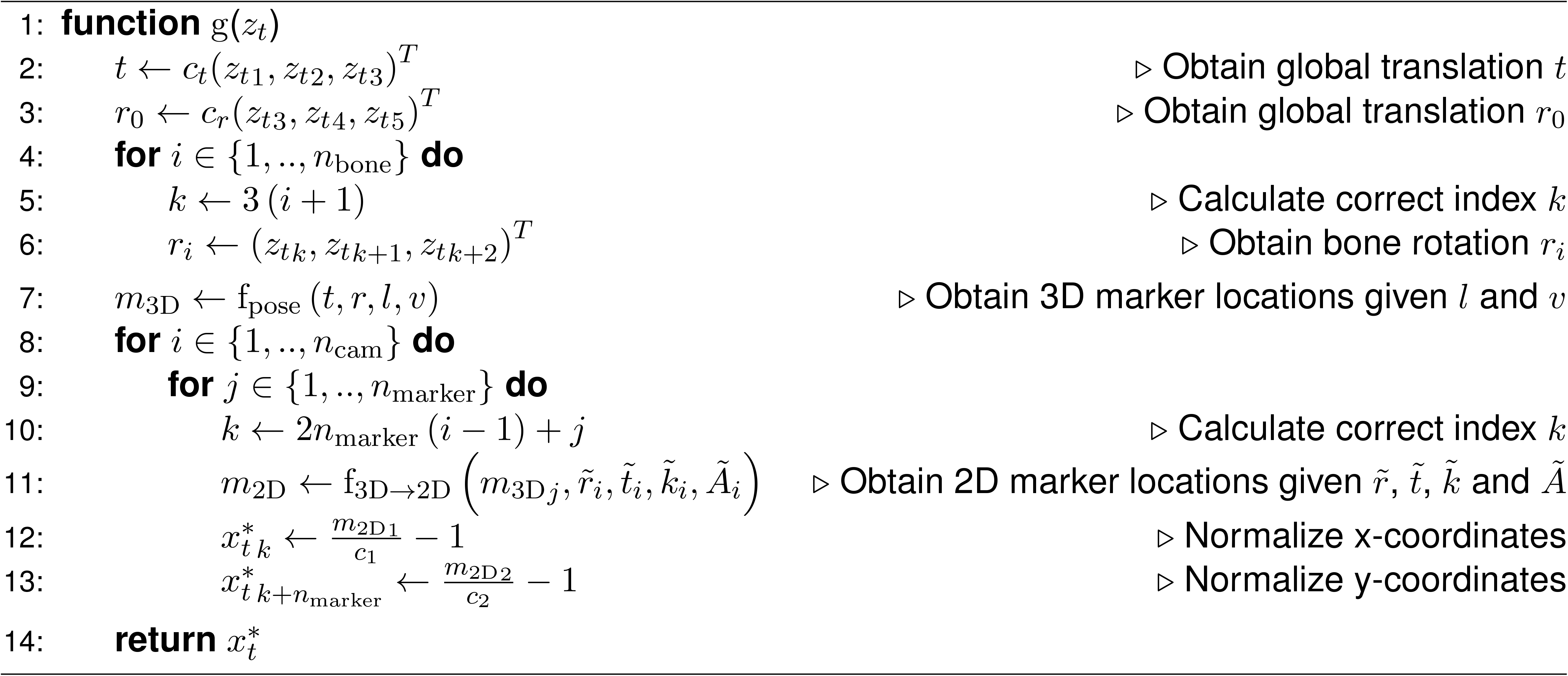

### 4.2 Theory of the expectation-maximization algorithm

While the EM algorithm was first introduced by Dempster et al. [5], we follow the concepts and notations stated by Bishop [1] and Murphy [9]. To derive a formulation of the ELBO we first note that the model’s joint distribution p (*x, z*) is equal to the product of the model’s likelihood p (*x* | *z*) and prior p (*z*):

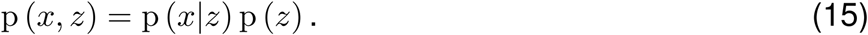

Additionally, we also note that the mutual dependency of the model’s marginal likelihood p (*x*), posterior p (*z*|*x*), likelihood p (*x*|*z*) and prior p (*z*) is given by Bayes’ theorem:

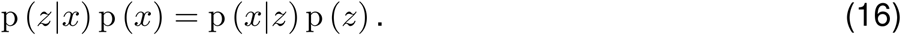

We now define an arbitrary probability density function q (*z*) over our state variables *z*, for which we know the following statement is true by definition:

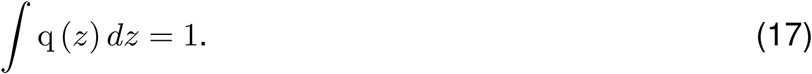

Multiplying equation 17 with an arbitrary constant *c* yields:

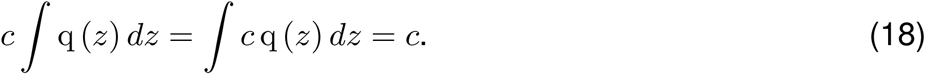

We can now replace the constant *c* with a function independent from the state variables *z* without loss of generality. If we choose this function to be the model’s marginal log-likelihood ln p (*x*), we obtain:

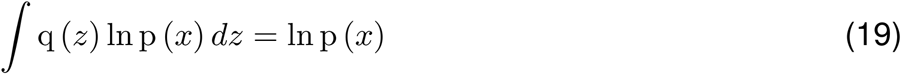

and note that, due to equation 17 and 18 respectively, the marginal log-likelihood ln p (*x*) is actually independent of the probability density function q (*z*). Next, we can use equation 15 and 16 to derive a relationship between the marginal log-likelihood ln p (*x*), the Kullback–Leibler (KL) divergence KL (*q*||*p*) and the ELBO ℒ, starting from equation 19:

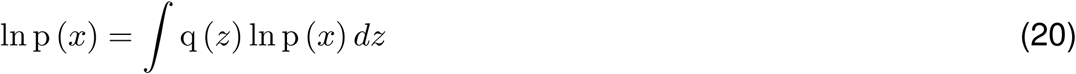

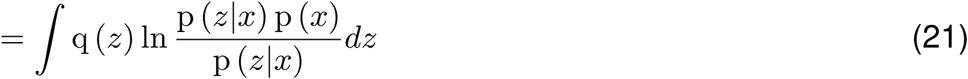

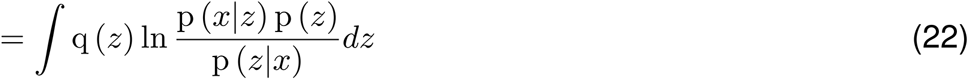

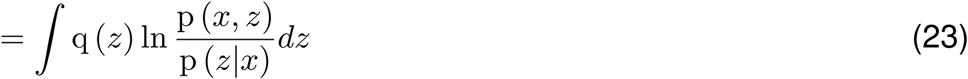

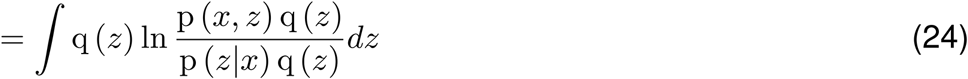

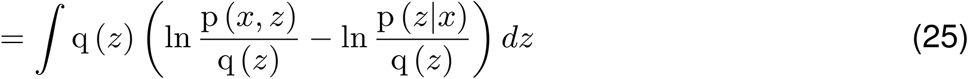

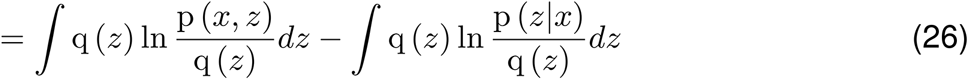

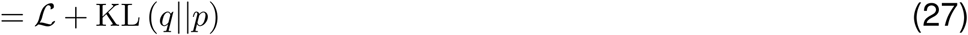

with 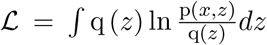 and 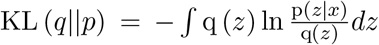. The KL divergence is a distance measure between the probability density functions q and p and as such always larger or equal to zero:

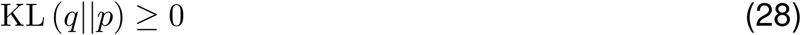

with equality KL (*q*||*p*) = 0 if *q* = *p*. When we add the ELBO 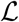 to equation 28 and combine the result with the derived definition of ln p (*x*), it becomes clear that the ELBO 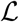 is a lower bound of the marginal log-likelihood:

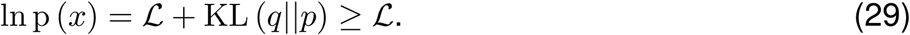

If we now acknowledge that we also require the model parameters Θ to compute the above quantities, i.e.

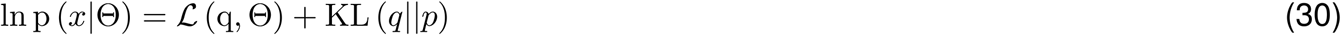

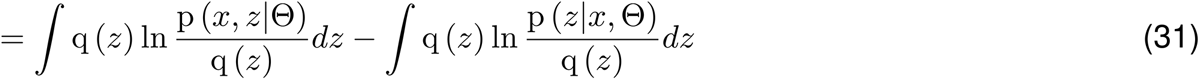

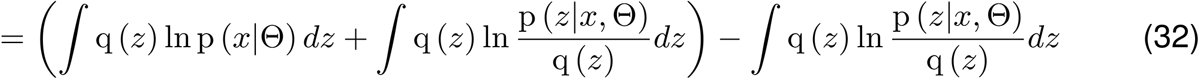

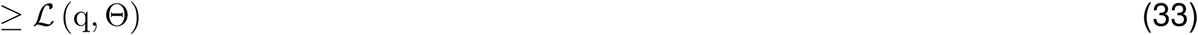

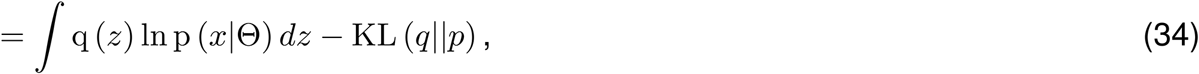

we can start building an understanding for how the EM algorithm works. In the E-step we are holding Θ constant and maximize 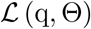 with respect to q, i.e. given a current estimate for the model parameters Θ_*k*_ we infer the probability density functions of our state variables p (*z*|*x*, Θ_*k*_), such that q (*z*) = p (*z*|*x*, Θ_*k*_), making the KL divergence KL (*q*||*p*) become zero, i.e. KL (*q*||*p*) = KL (*p*||*p*) = 0, and the marginal log-likelihood ln p (*x*|Θ_*k*_) become equal to the ELBO 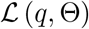. Here, setting q (*z*) = p (*z*|*x*, Θ_*k*_) maximizes the ELBO 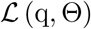 due to the equality given by equation 34 and the previously mentioned fact that the marginal log-likelihood ln p (*x*) is actually independent of the probability density function q (*z*). Subsequently, in the M-step we are holding q constant and maximize 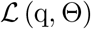 with respect to Θ in order to obtain a new set of model parameters Θ_*k*+1_, leading to an increased marginal log-likelihood ln p (*x*|Θ_*k*+1_), as the KL divergence becomes greater then zero again, i.e. KL (*q*||*p*) ≥ 0 and 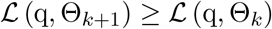. Thus, the starting point in the M-step is the following:

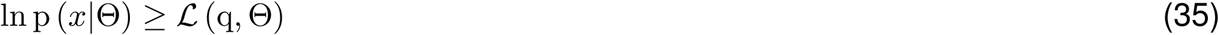

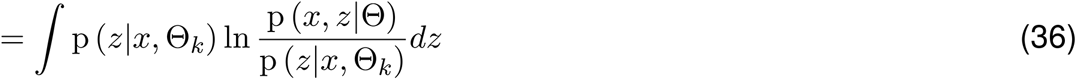

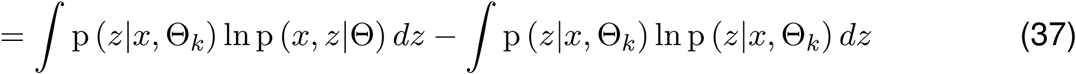

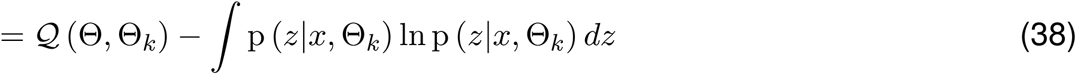

with 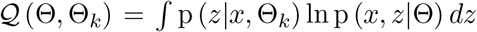. We note that the latter term is independent of Θ and can be omitted since our goal is to optimize the ELBO 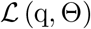 with respect to Θ. Therefore, instead of maximizing the ELBO 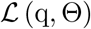 directly, we can just maximize the function 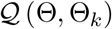. We furthermore notice that 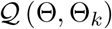 has the form of an expectation value, i.e. we can obtain 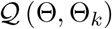 by taking the expectation of ln p (*x*, *z*|Θ) with respect to *z*:

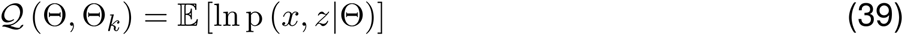

where 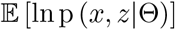 is conditioned on *x* and Θ_*k*_, i.e. both quantities are given. With this we finally arrive at the essence of what is done during the M-step, i.e. maximizing 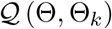 with respect to Θ to obtain new model parameters Θ_*k*+1_:

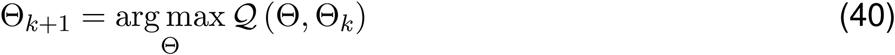

### 4.3 The unscented transformation

We are required to approximate expectation values to perform the E-step, i.e. when applying the unscented Kalman filter and the unscented Kalman smoother (Algorithm 7 and 9), as well as the M-step, i.e. when maximizing 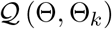 (equation 39), as we can not compute them analytically [8].

These expectation values are of the form:

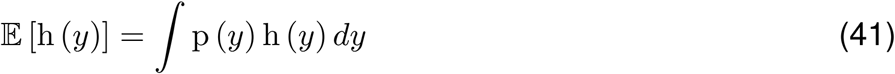

where h is an arbitrary function and *y* ∈ ℝ^*d*^ an arbitrary normally distributed random variable, i.e. 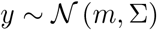. We can obtain such approximations using the unscented transformation f_ut_, which was first introduced by Julier et al. [7] and is defined in Algorithm 5. Given the mean *m* and the covariance Σ, the unscented transformation f_ut_ generates so called sigma points 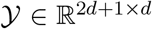, whose locations are systematically spread around the mean *m* based on the covariance Σ:

**Algorithm 5.**
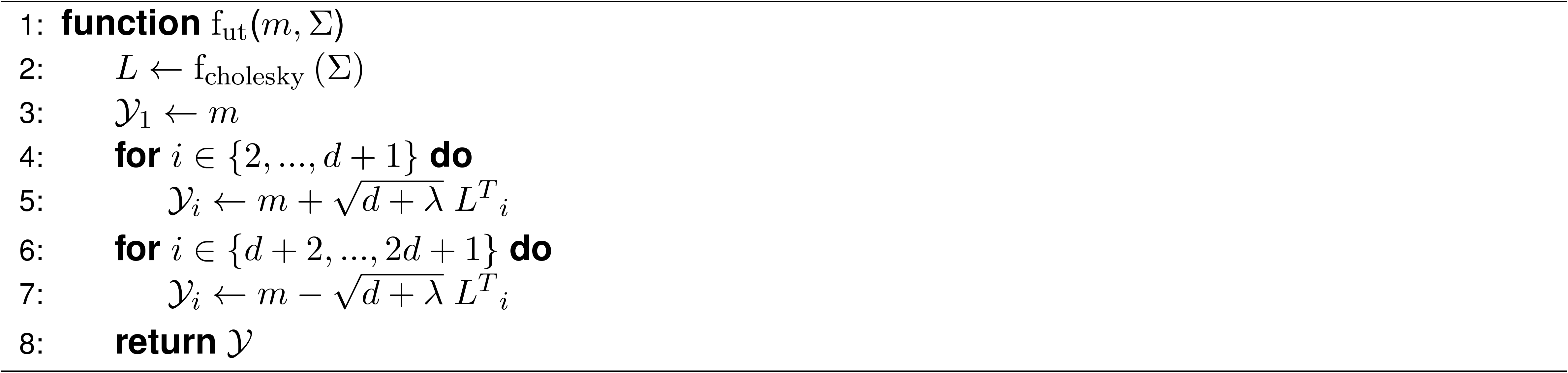

Here f_cholesky_ (Σ) denotes the Cholesky decomposition of matrix Σ, which computes a lower triangular matrix *L* such that *LL^T^* = Σ, and *λ* can be calculated as follows:

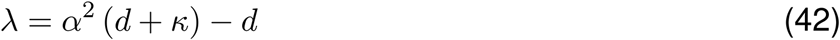

where we set the parameters *α* = 1 and *κ* = 0 [8], such that *λ* = 0. Using the sigma points 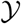, we can approximate 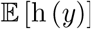 as follows:

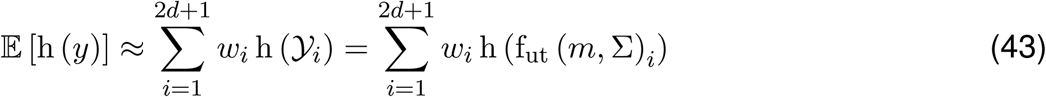

with the weights *w*:

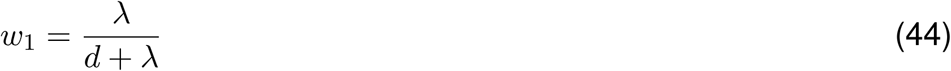

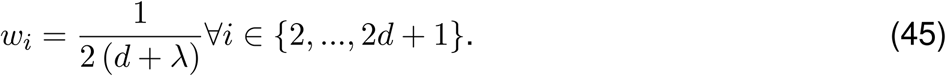

which due to our choice of *α* and *κ* simplifies to:

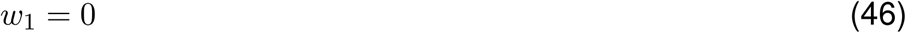

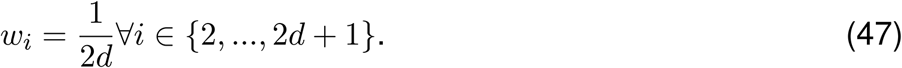

### 4.4 Expectation step

In the E-step we need to infer the probability density function of the latent variable *z_t_* for all time points *t* of a behavioral sequence, given the set of all measurements *x* and the model parameters Θ, noted as p (*z*_*t*_|*x*, Θ). Since all random variables of the model are assumed to be normally distributed, this property is maintained for the latent variable *z_t_* as well. Therefore, *z_t_* is drawn form a normal distribution with mean *μ_t_* and covariance *V_t_*, i.e. 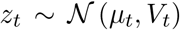. By using all measurements *x* of the sequence for the inference of p (*z*_*t*_|*x*, Θ) at time point *t*, information of the past as well as of the future is processed, which is what the unscented RTS smoother is used for. However, to derive the equations of the smoother we first need to focus on the inference when only information of the past is available, i.e. we want to infer p (*z_t_*|*x*_1_, …, *x_t_*, Θ) where only measurements until time point *t* are given, which can be achieved by utilizing the unscented Kalman filter. To avoid confusions, we denote mean values and covariance matrices obtained from the unscented Kalman smoother as 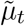 and 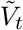, whereas those calculated via the unscented RTS smoother are denoted as 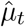 and 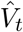.

#### 4.4.1 The unscented Kalman filter

The unscented Kalman filter is an iterative algorithm, which calculates the filtered values for the mean 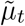 and covariance 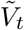 at a time point *t*, based on the filter output for these values 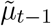 and 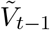 at the previous time point *t* − 1 as well as the measurement variable *x_t_* for time point *t*. The inference scheme for obtaining p (*z_t_*|*x*_1_, …, *x*_*t*−1_, Θ) is given by Algorithm 6 and 7 [11, 12]:

**Algorithm 6.**
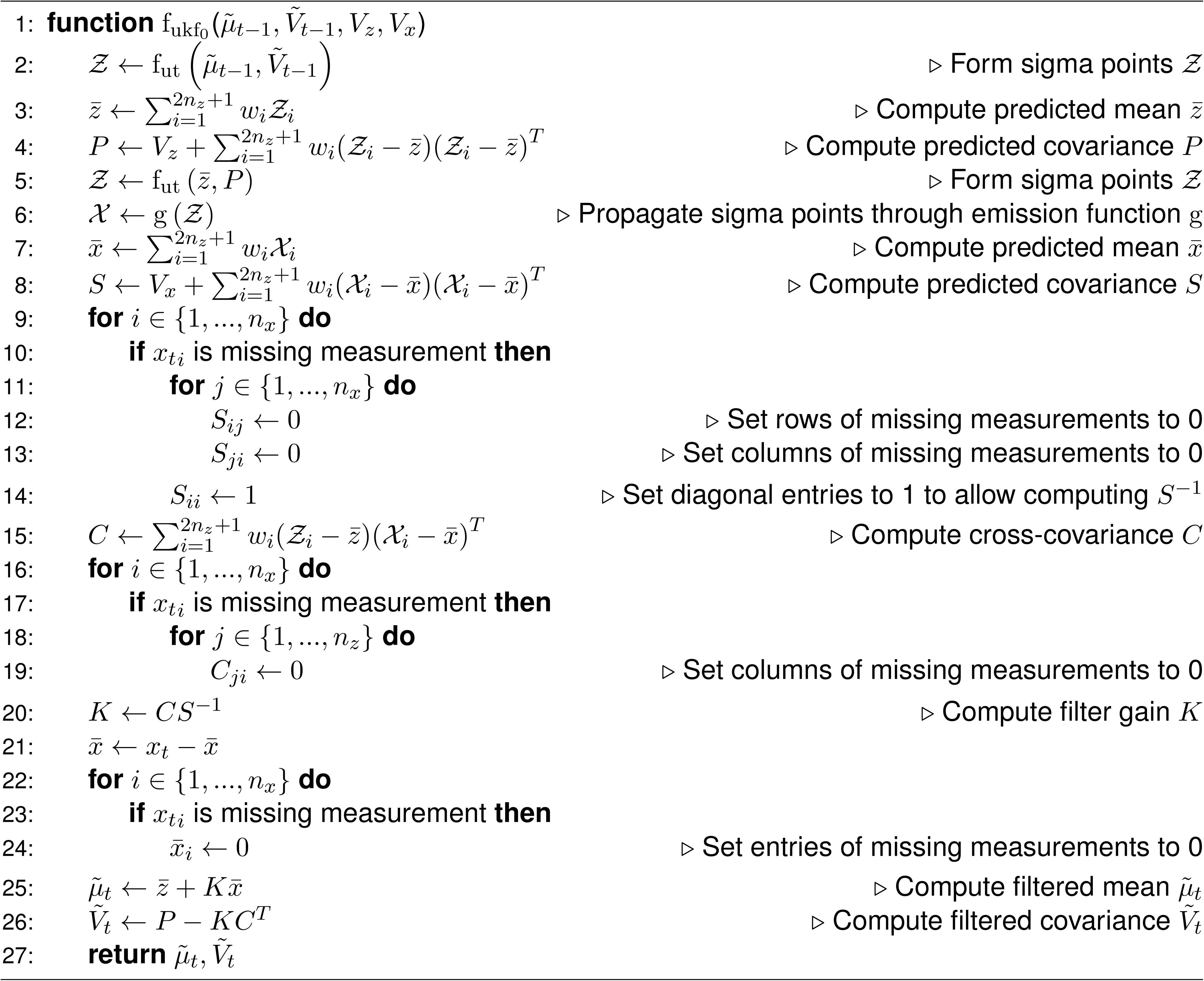

To obtain values for filtered means 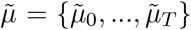 and covariances 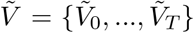 for all time points one needs to iterate through the entire behavioral sequence:

**Algorithm 7.**
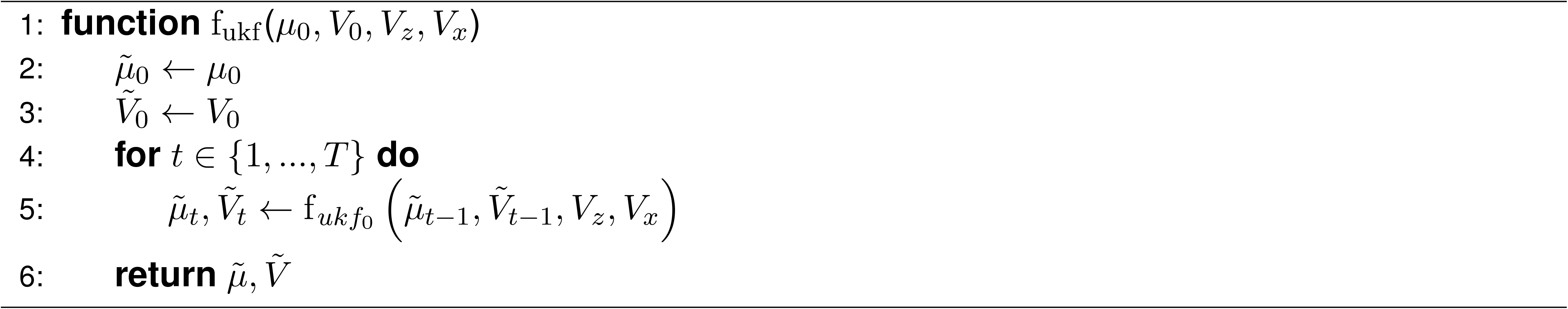

#### 4.4.2 The unscented RTS smoother

The unscented RTS smoother is also an iterative algorithm, which calculates the smoothed values for the mean 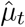 and covariance 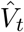 at a time point *t*, based on the smoother output for these values 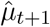 and 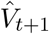 at the next time point *t* + 1 as well as the corresponding output from the unscented Kalman filter 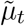 and 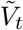 for time point *t*. The inference scheme for obtaining p (*z_t_*|*x*, Θ) is given by Algorithm 8 and 9 [12]:

**Algorithm 8.**
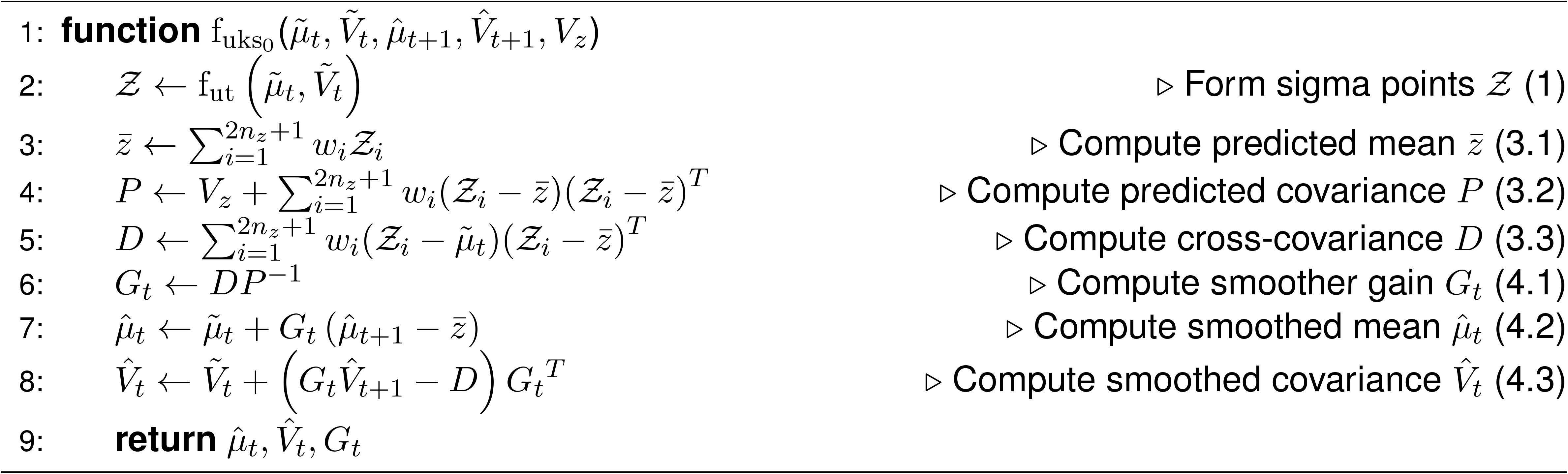

To obtain values of the smoothed means 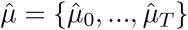 and covariances 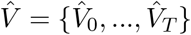 for all time points one needs to run the forward filtering path and then iterate backwards through the entire behavioral sequence:

**Algorithm 9.**
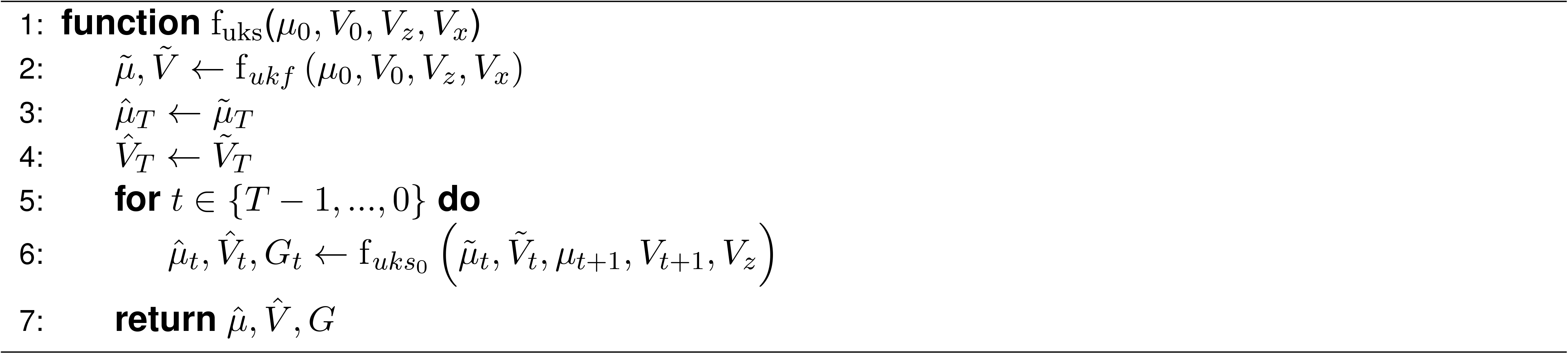

Here, the set of all smoother gains *G* = {*G*_0_, …, *G*_*T*−1_} is needed for performing the M-step later on.

#### 4.4.3 Enforcing anatomical constraints

The plain formulation of the unscented RTS smoother does not allow constraining the state variables. However, in order to enforce joint angle limits we need to ensure that Rodrigues vectors encoding bone rotations stay within specified limits. Therefore, we introduce a mapping function 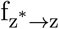, which allows for mapping a redefined state variable 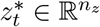 onto the original one 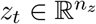, while enforcing that entries of *z_t_* corresponding to bone rotations stay within their respective lower and upper bounds. The respective mapping function 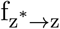 is given by Algorithm 10.

**Algorithm 10.**
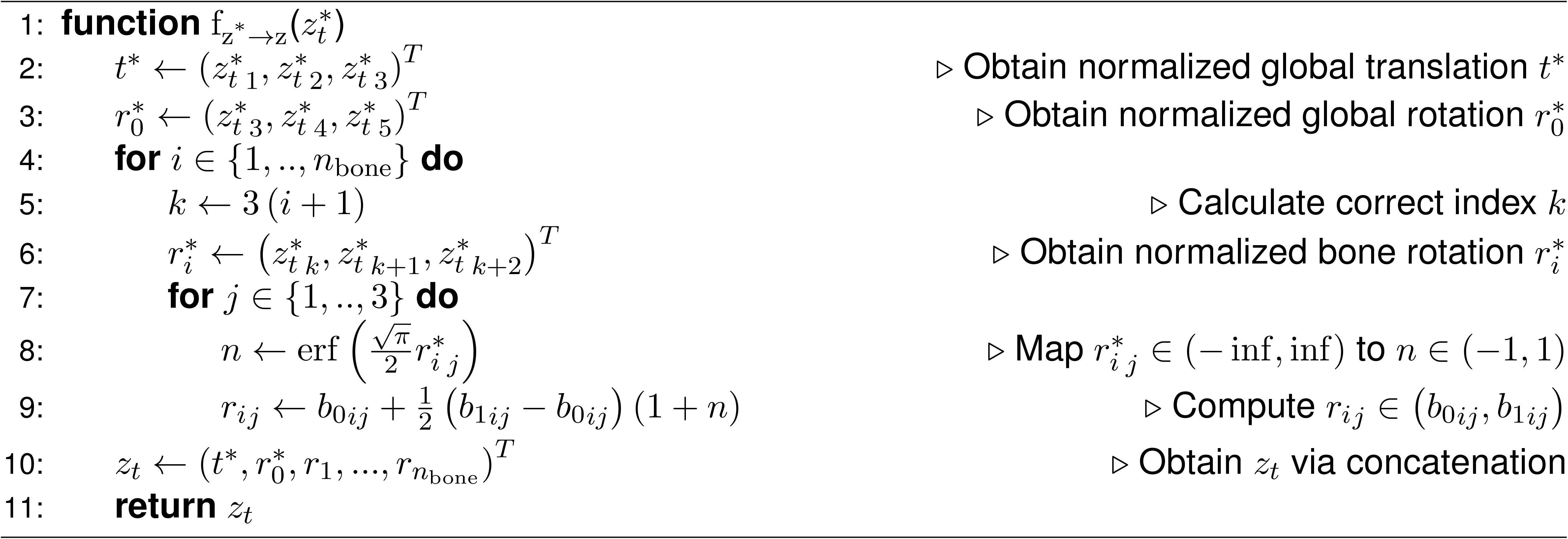

Here, *b*_0*ij*_ and *b*_1*ij*_ denote the lower and upper bound corresponding to entry *j* of the Rodrigues vector *r_i_*, which encodes the rotation of bone *i*, and erf is a sigmoidal function, i.e. the error function given by:

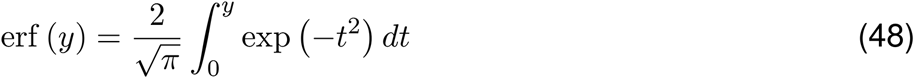

for *y* ∈ ℝ. In order to enforce joint angle limits we just replace the original transmission and emission equation in our state space model given by equations 13 and 14 with:

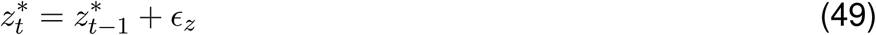

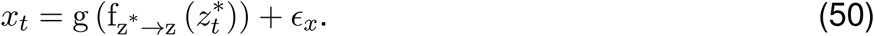

In the following we always refer to the state space model given by equations 49 and 50 and therefore also to the redefined state variables *z*^∗^ but we drop ∗ in the notation.

### 4.5 Maximization step

In the M-step we find a new set of model parameters Θ_*k*+1_ by maximizing the ELBO 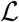, given the smoothed means 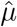 and covariances 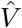 as well as the smoother gains *G*, which we obtained in the E-step using a current estimate of the model parameters Θ_*k*_.

#### 4.5.1 Obtaining new model parameters by maximizing the evidence lower bound

We can take advantage of the specific structure of the state space model when maximizing the ELBO 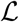 [8]. In the state space model the state variables fulfill the Markov property, i.e. each state variable *z_t_* only depends on the previous one *z*_*t*−1_. Based on this we can compute the model’s joint distribution:

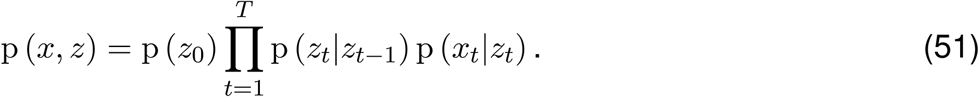

When we now take the logarithm of the joint distribution and acknowledge that the model parameters Θ are also required for computing the joint distribution we obtain:

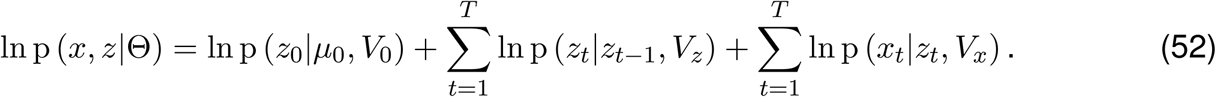

However, to maximize 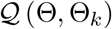 we actually need to consider the expectation value of ln p (*x*, *z*|Θ):

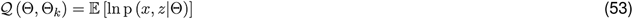

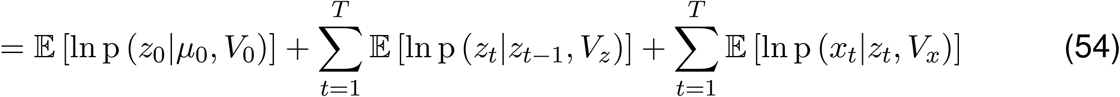

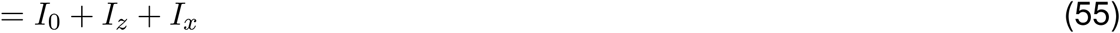

with 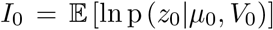, 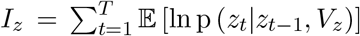 and 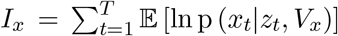. If we now acknowledge that all random variables in our state space model are normally distributed, i.e. 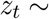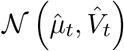, it becomes clear that computing 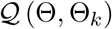 only involves evaluating the expectation values of log-transformed normal distributions (see Appendix A). Consequently, we can obtain simplified terms for the individual components *I*_0_, *I_z_* and *I_x_* of 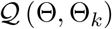 using the smoothed means 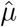 and covariances 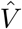 as well as the smoother gains *G*. For *I*_0_ we get:

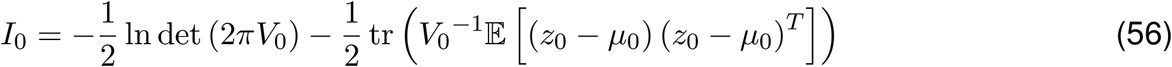

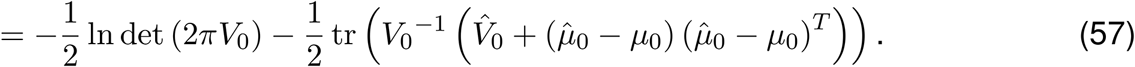

To obtain a simplified expression for *I_z_* we need to form pairwise sigma points 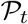, (as there are always two random variables involved simultaneously, 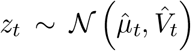 and 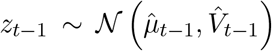, when evaluating the expectation values of the underlying log-transformed normal distributions in *I_z_*. For each of the *T* transition steps we generate the pairwise mean vector 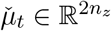:

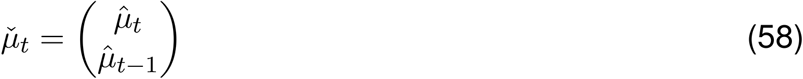

as well as the pairwise covariance matrix 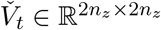:

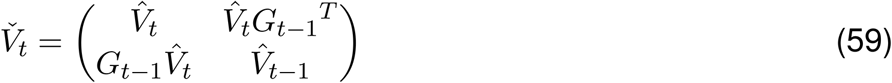

and calculate the pairwise sigma points 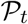 as follows:

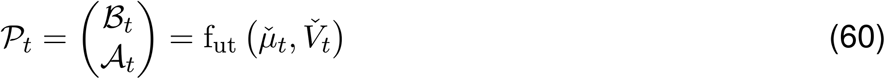

where concatenating the incomplete pairwise sigma points 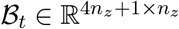 and 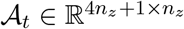 gives 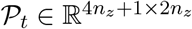. Consequently, the weights 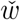 associated with the pairwise sigma points 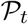 are then given in accordance with the concepts discussed in Section 4.3:

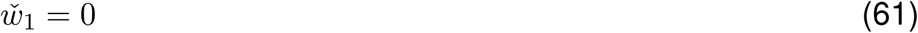

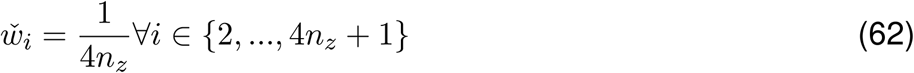

A simplified term for *I_z_* is then given by:

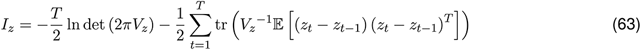

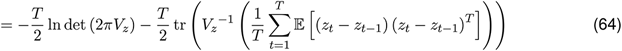

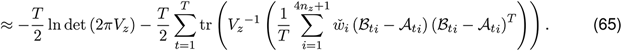

To evaluate the expectation value in *I_x_* it is sufficient to just use the normal sigma points 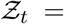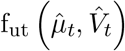 and propagate them through our emission function g:

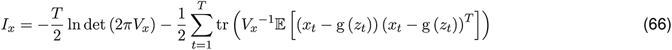

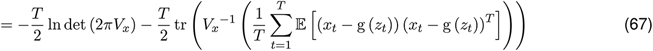

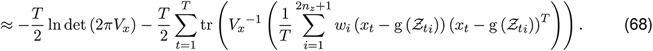

To finally obtain new model parameters 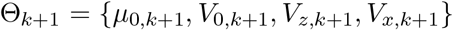 we still need to differentiate 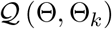 with respect to *μ*_0_, *V*_0_, *V_z_* and *V_x_*, set the resulting derivatives to zero and solve them for *μ*_0_, *V*_0_, *V_z_* and *V_x_* respectively. The required derivatives have the following form (see Appendix B):

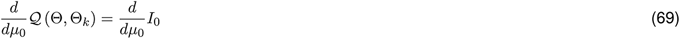

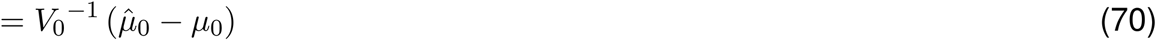

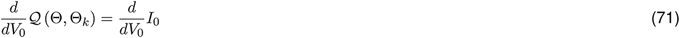

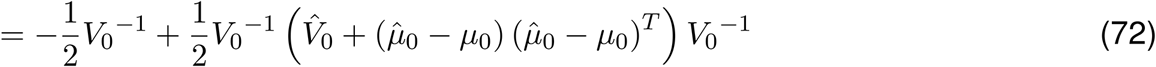

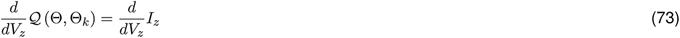

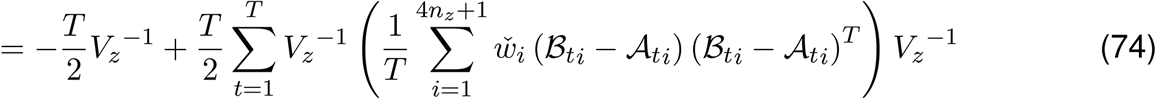

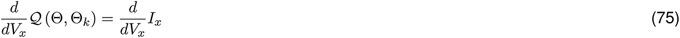

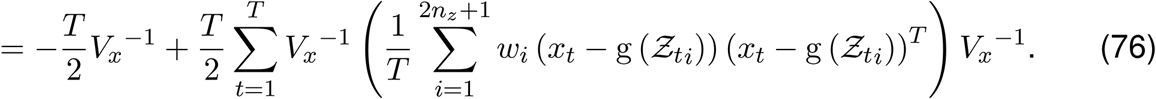

Setting these derivatives to zero and solving for *μ*_0_, *V*_0_, *V_z_* and *V_x_* yields the following:

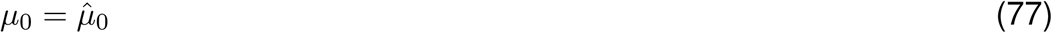

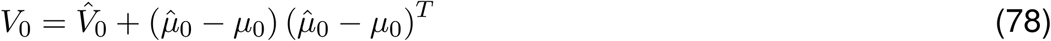

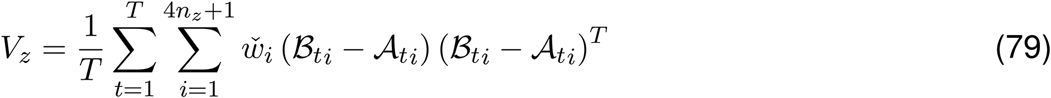

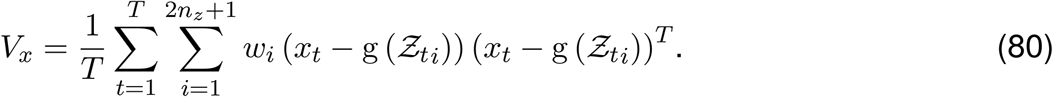

The resulting values for *μ*_0,*k*+1_, *V*_*z,k*+1_ and *V*_*x,k*+1_ are then given by equations 77, 79 and 80. To obtain *V*_0,*k*+1_ we need to substitute *μ*_0,*k*+1_ into equation 78, giving 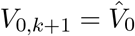. Lastly, we still need to adjust the solution for *V*_*x,k*+1_ to also account for missing measurements. Besides that, we note that t is sufficient to only compute the diagonal entries of *V*_*x,k*+1_, since we enforce the covariance matrix of the measurement noise *V_x_* to be a diagonal matrix. Thus, the final solution for a diagonal entry *j* ∈ {1, …, *n_x_*} of *V*_*x,k*+1_ is given by:

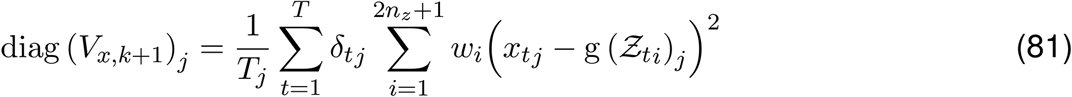

where the function diag gives the diagonal entries of the input matrix, *δ_tj_* indicates if at time point *t* the entry *j* of *x_t_* is associated with a missing measurement, i.e. *δ_tj_* = 0, or not, i.e. *δ_tj_* = 1, and *T_j_* is the total number of successful measurements for entry *j* in the entire behavioral sequence, i.e. 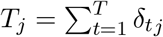.

### 4.6 Convergence of the expectation-maximization algorithm

We calculate the changes in the model parameters Θ in each iteration *k* of the EM algorithm to check for convergence [1]. Particularly, we are computing the vectors 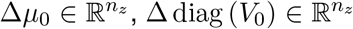, 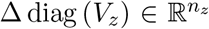 and 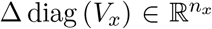, which contain the relative changes of *μ*_0_, *V*_0_, *V_z_* and *V_x_* at iteration *k*:

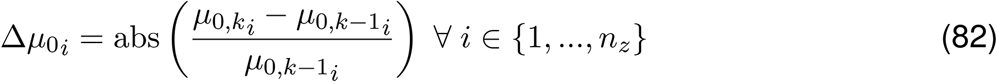

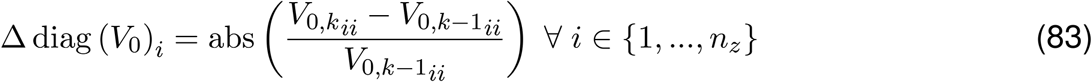

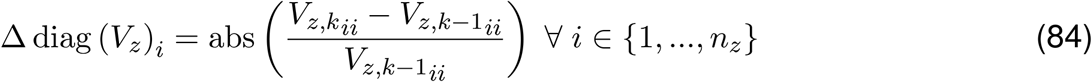

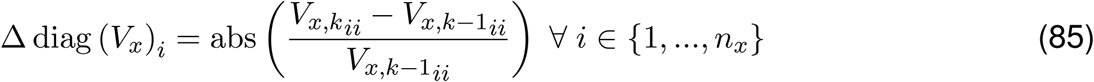

where abs is a function returning the absolute value of its input argument and *μ*_0,*k*_, *V*_0,*k*_, *V_z,k_* and *V_x,k_* are the model parameters at iteration *k* whereas *μ*_0,*k*−1_, *V*_0,*k*−1_, *V_z,k_*_−1_ and *V_x,k_*_−1_ are those at iteration *k* − 1. We only focus on the diagonal entries of the covariances *V*_0_ and *V_z_* since a fraction of their off-diagonal entries is expected to be zero. Using these relative changes we construct a vector 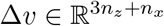 containing all relative changes via concatenation:

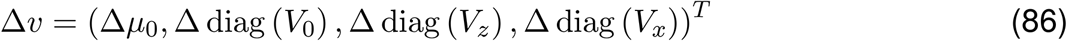

and assume convergence is reached when the mean 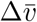 of Δ*v* falls below a threshold *∊*_tol_:

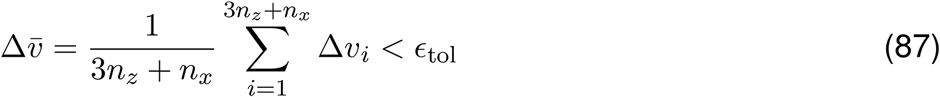

where we set *∊*_tol_ = 0.05.

### 4.7 Implementation of the expectation-maximization algorithm

We initialize the mean of the state variables *μ*_0_ by minimizing the objective function given by equation 9 but keep the bone lengths *l* and the surface marker positions *v* constant and set *n*_time_ = 1, i.e. we only include a single time point in the optimization, which is identical to the first time point of a respective behavioral sequence. The covariances *V*_0_, *V_x_* and *V_z_* are initialized as matrices whose diagonal elements all equal 0.001 and off-diagonal entries are set to zero. To learn new model parameters *μ*_0_, *V*_0_, *V_x_* and *V_z_* we run the EM algorithm, given by Algorithm 11, with the stated initial values using measurements *x* obtained from the behavioral sequence. Finally, once the EM algorithm converged, we use the unscented RTS smoother with the resulting learned model parameters to reconstruct poses of the behavioral sequence.

**Algorithm 11.**
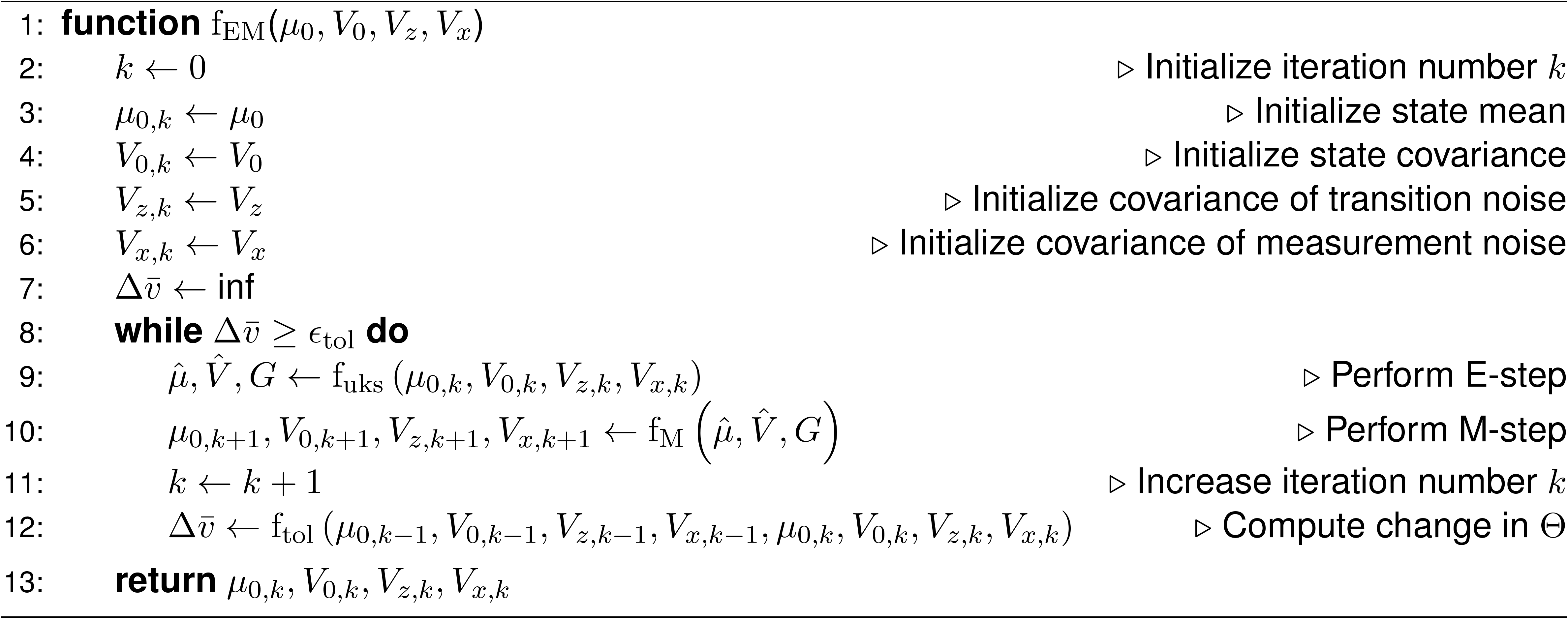

Here, in accordance to the concepts stated in Section 4.5 and 4.6, function f_M_, given by Algorithm 12, performs the M-step and function f_tol_, given by Algorithm 13, computes the mean 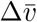 of the relative changes of the model parameters Θ.

**Algorithm 12.**
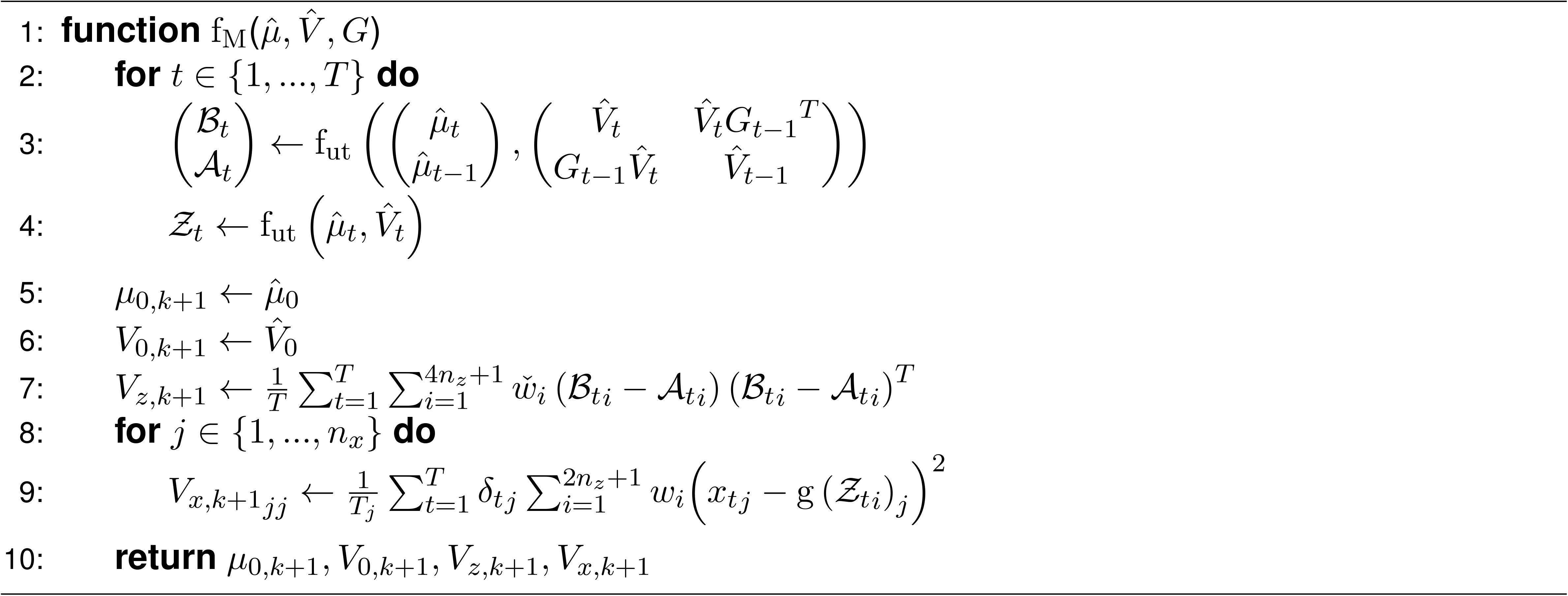

**Algorithm 13.**
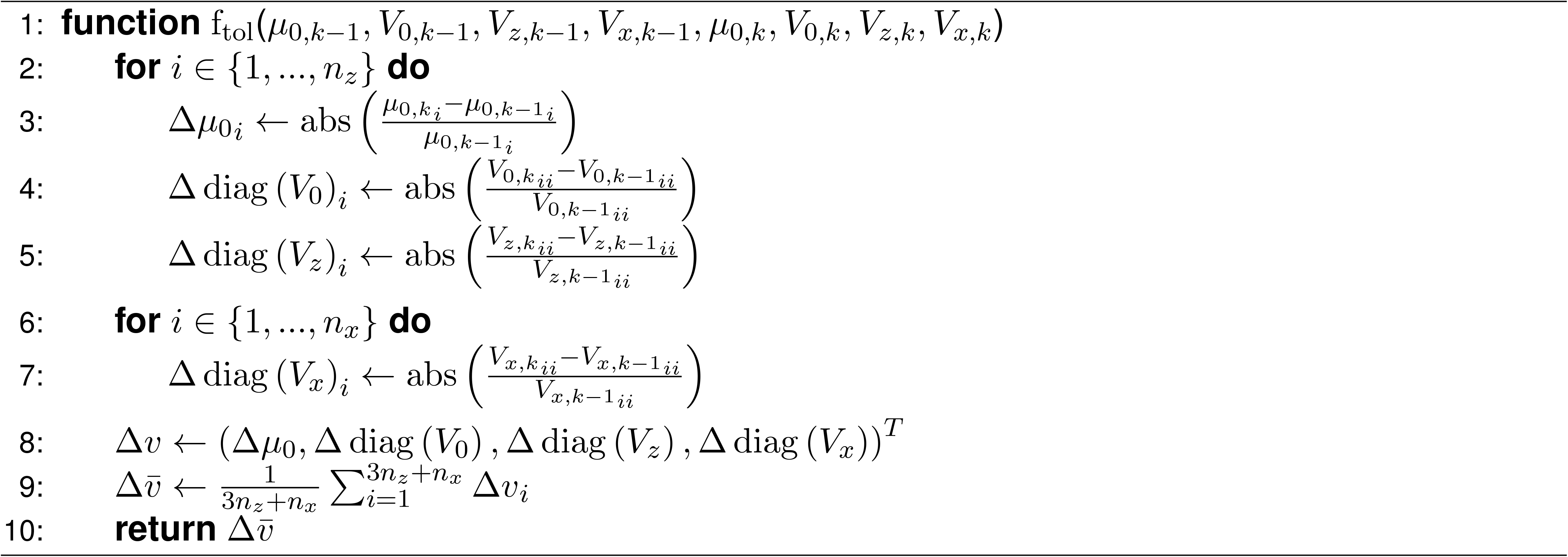

## Appendix

### A Evaluating expected values of log-transformed normal distributions

Given a *d*-dimensional normal distribution p_norm_ with mean *μ_y_* ∈ ℝ^*d*^ and covariance *V_y_* ∈ ℝ^*d*×*d*^, evaluating it for a normally distributed random variable 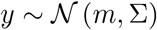 takes the following form:

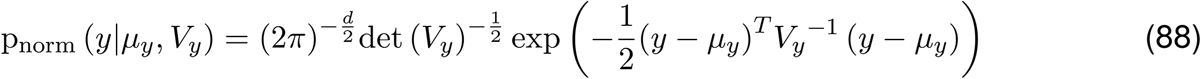

where det (*V_y_*) ∈ ℝ denotes the determinant of matrix *V_y_*. Applying a logarithmic transformation yields:

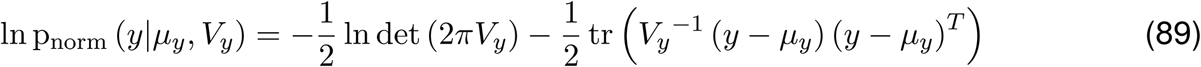

where tr (*V_y_*) ∈ ℝ denotes the trace of matrix *V_y_*. Noticing that 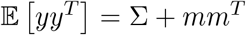 [8], we can take the expected value of equation 89 with respect to *y* and obtain:

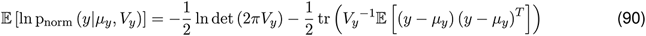

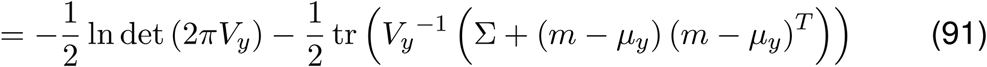

### B Derivatives

Given a *d*-dimensional vector *v* ∈ ℝ^*d*^, two symmetric matrices *M* ∈ ℝ^*d*×*d*^ and *C* ∈ ℝ^*d*×*d*^ as well as a scalar *c* ∈ ℝ, we can obtain the following derivatives:

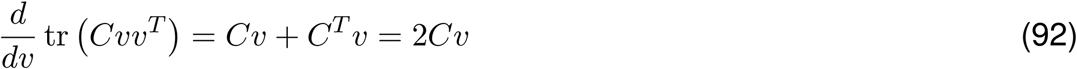

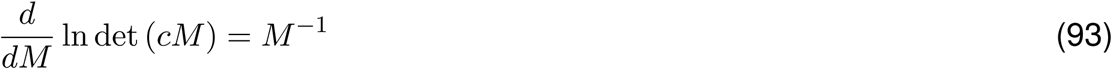

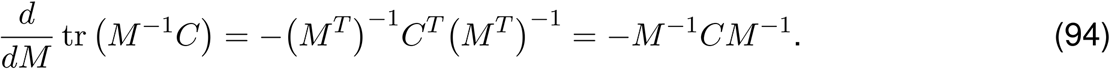

## Notes

### Competing Interest Statement

The authors have declared no competing interest.

### Summary of Updates

The Version of Record of this article is published in Nature Methods, and is available online at https://dx.doi.org/10.1038/s41592-022-01634-9

